# A genome-scale single cell CRISPRi map of *trans* gene regulation across human pluripotent stem cell lines

**DOI:** 10.1101/2024.11.28.625833

**Authors:** Claudia Feng, Elin Madli Peets, Yan Zhou, Luca Crepaldi, Sunay Usluer, Alistair Dunham, Jana M Braunger, Jing Su, Magdalena E Strauss, Daniele Muraro, Kimberly Ai Xian Cheam, Marc Jan Bonder, Edgar Garriga Nogales, Sarah Cooper, Andrew Bassett, Steven Leonard, Yong Gu, Bo Fussing, David Burke, Leopold Parts, Oliver Stegle, Britta Velten

## Abstract

Population-scale resources of genetic, molecular, and cellular information form the basis for understanding human genomes, charting the heritable basis of disease, and tracing the effects of mutations. Pooled perturbation assays applied to cellular models, probing the effect of many perturbations coupled with an scRNA-seq readout (Perturb-seq), are especially potent references for interpreting disease-linked mutations or gene expression changes. However, the utility of existing maps has been limited by the comprehensiveness of perturbations possible, and the relevance of their cell line context. Here, we present the first genome-scale CRISPR interference (CRISPRi) perturbation map with single-cell RNA sequencing readout across many human genetic backgrounds in human pluripotent cells. To do so, we establish large-scale CRISPRi screening in human induced pluripotent stem cells from healthy donors, using over 20,000 guide RNAs to target 7,226 genes across 34 cell lines from 26 genetic backgrounds, and gather expression data from nearly 2 million cells. We comprehensively map *trans* expression changes induced by the target knockdowns, which complement co-expression patterns in unperturbed cells and facilitate the functional annotation of target genes to biological processes and complexes. Consistency of targeting protein complex members point to protein complexes as a nexus for aggregating transcriptional variation, revealing novel interaction partners. We characterise variation in perturbation effects across donors, with expression quantitative trait loci linked to higher genetic modulation of perturbation effects but overall low replication of trans effects due to knockdown of their corresponding cis regulators. This study pioneers population-scale CRISPR perturbations with single cell readouts that will fuel foundation models for the future of effective modulation of cellular disease phenotypes.

## Introduction

Cellular models allow interrogating disease phenotypes and basic processes in controlled experiments. Undifferentiated induced pluripotent stem cells (iPSCs) are an established system for modelling human development and disease^1–3^. These cells are generated by transforming easy-to-acquire cell types, such as human fibroblasts, into an embryonic-like state, where cells have the capacity to differentiate into the three germ layers. Given the cell-type specificity of many diseases and their inherent ability to self-renew and differentiate, iPSCs represent a powerful tool for studying human variation in cell types that are otherwise difficult to obtain. Consequently, many efforts have been made in identifying the effects of common variation on molecular phenotypes in these cells, e.g. expression quantitative trait locus (eQTL) of common, rare and structural variants^1–3^. While this identified thousands of loci altering expression levels of nearby genes in *cis*, the regulatory landscape underlying the downstream consequences on cellular pathways and function are to a large extent still poorly understood. In particular, existing studies on *trans* effects, in particular in human pluripotent cells, are underpowered due to insufficient genomic resources and small effect sizes of genetic variation in the natural population^3^.

To complement natural genetic variation, CRISPR has recently emerged as a powerful tool for gene editing, silencing or activation in a targeted and cost-efficient manner. By combining pooled CRISPR-based screening with single-cell gene expression as a read-out, we are now able to study molecular consequences of genetic perturbations on candidate genes^4–8^, identify their downstream targets and infer their biological function and relation to disease. Where non-interventional single-cell expression studies can identify co-expressed genes, inducing a genomic perturbation provides directionality on the nature of a regulatory relationship. To date, CRISPR-based screens have mainly been used to elucidate gene function^9^ and to identify regulatory networks^5,10^. However, resources that map perturbation responses at a genome-scale remain scarce and despite their relevance for understanding human disease and development comprehensive screens have not yet been conducted in iPSCs. Further, while it is well-known that genetic background matters both for mutation impact on disease risk^11^, as well as its effect in a functional assay^1,3,12^, no efforts have been made to account for different genetic backgrounds in existing studies so far^13–15^ .

Here, we present results from a genome-scale CRISPRi screen conducted in iPSCs derived from tens of individuals with scRNA-seq readout. By combining two distinct experimental designs, leveraging the number of perturbations in our dataset, as well as a diverse set of genetic backgrounds, we show that this population genomic perturbation approach with single cell phenotyping can be used to recover known regulatory networks, study cell type-specific biology and compare effects of synthetic dosage change and natural genetic variation. This work represents the second genome-scale CRISPRi screen with scRNA-seq readout and is the only study that accounts for the role of genetic background on gene function, establishing a necessary and fundamental building block for conducting population-scale genetic analyses using genome perturbation techniques.

## Results

### Measuring knockdown effects across multiple donors at scale

First, we set out to chart the landscape of genetic perturbation effects in multiple individuals by performing gene knockdowns using CRISPRi coupled with a single-cell RNA-seq readout (**Figure 1A, Methods**). To do so, we targeted 7,226 genes in 34 high-quality iPS cell lines derived from 26 distinct healthy donors. The targets were selected to cover genes whose knockdown in iPSCs or cancer cell lines^16^ exhibited growth defects (2,264 genes and 4,594 genes, respectively, of which 1,725 genes exhibited fitness effects in both) or are highly expressed in iPSCs (2,093 additional genes; **Supplementary Table ST1-1**, **Figure S1A, Methods**). The targeting library included three guide RNAs (gRNAs) per gene from the Dolcetto library^17^, as well as 40 non-targeting guides. Following quality control, demultiplexing of cell lines from genotypes, and gRNA assignment (**Methods**) we obtained 219,206 cells assigned to a source cell line and a targeting guide (median of 8 cells per gRNA and 25 cells per gene; **Methods**, **Figure S1B, Supplementary Table ST1-2**), as well as 499,998 control cells assigned to a donor, but either no guide or a non-targeting gRNA. We considered target genes with a minimum coverage of 10 cells (6,673/7,226, 92%) for further analysis based on power calculations (**Figure S1C**, **Methods**), and computed log-fold changes that quantify the average perturbation effects across all cell lines for each target (6,673 targets x 6,471 expressed genes; **Figure 1D**, **Methods**).

**Figure 1.**
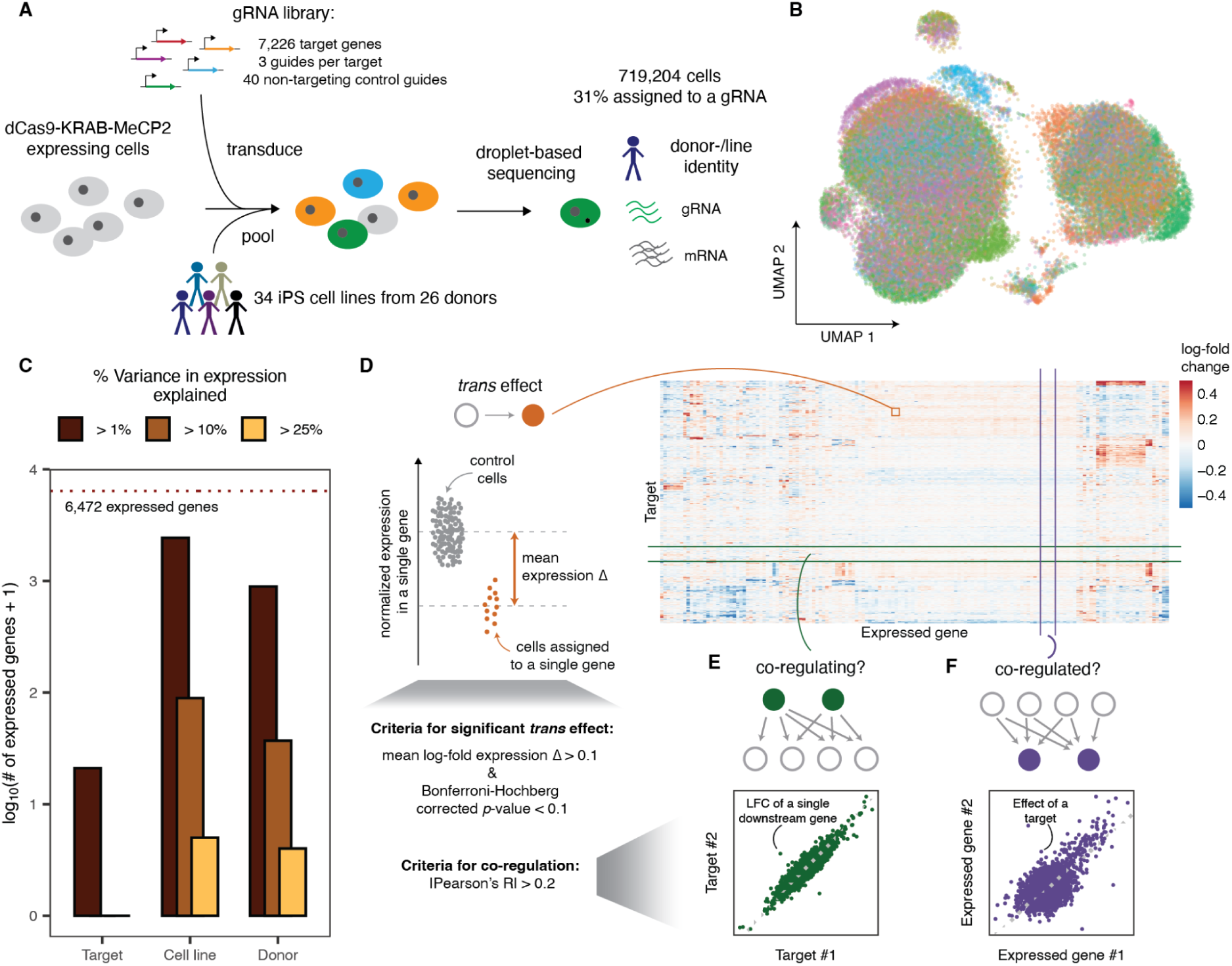
| Quantifying knockdown effects in cells from multiple donors at scale. A) Experimental design of a genome-scale CRISPRi screen with single-cell RNA-seq read-out from 34 human iPSC cell lines. B) UMAP of the expression profiles of cells assigned to a donor and targeting guide after correcting for technical covariates, colored by cell line. C) Variance decomposition, displaying relative effects of cell line, genomic background and target gene perturbation on gene expression. D) Overview of the analysis strategy to understand individual *trans* effects (orange), E) similarity of effects across all genes (green), F) and similarity of perturbation responses across all targets (purple).

We next quantified the sources of global variation in the gene expression profiles. As expected for a large-scale single cell RNA sequencing study, technical factors, cell quality and cell line were major drivers of expression heterogeneity between cells (**Figure 1B-C, S1D**). Transcriptional changes induced by CRISPRi knockdown were relatively small compared to those global sources of heterogeneity (**Figure 1C**), suggesting that CRISPRi targeting induces mostly subtle transcriptional changes rather than complete cell state shifts within the considered time period of 3-6 days post-infection (DPI).

The quality of CRISPR targeting can be evaluated based on the on-target expression changes, as well as its effect on downstream genes beyond the target gene (“*trans effect*”). The mean expression of a targeted gene was significantly lower (Benjamini-adjusted p-value < 0.1) compared to control cells for 2,900 of the 4,874 (60%) knockdowns targeting an expressed gene (median log-fold change = -0.42 across all significant on-target *trans* effects), confirming an overall efficient target down-regulation in our system. In contrast, only 0.02% of all target-expressed gene combinations showed evidence of downregulation in *trans* beyond the target (8,833 out of 43,176,109 tested target-expressed gene combinations, **Figure S2-1A**). To assess the quality of our measured *trans* effects, we considered a set of well characterised transcription factors with known regulatory relationships^18^ across diverse cell types, states, and transitions. All of the 17 significant *trans* effects in our data that overlapped these annotations (**Figure S2-1B**) were concordant with the effect direction expected from the function of the transcription factor. Specifically, knockdown of known activators (16 of 17 significant *trans* effects) resulted in reduced expression of the downstream genes, while knockdown of the known repressor *BACH1* caused up-regulation of its downstream target, *HMOX1*. This remarkable concordance (p = 0.02, binomial test), highlights the potential for validating and charting gene regulatory effects from our data.

### Global transcriptional changes caused by gene-dosage reduction

Signatures of perturbation effects can reflect regulatory mechanisms in three ways: (i) a *trans* effect of a target gene on downstream genes reflects links in a regulatory network (**Figure 1D**), (ii) correlation of *trans* effects for two target genes indicates similar function of the targets (**Figure 1E**), and (iii) correlation of gene responses in *trans* indicates common control by shared regulators (**Figure 1F**). We next elucidated the factors that determine gene expression control in *trans*, shared *trans* effects (co-regulating targets), and shared regulators (co-regulated expressed genes).

Perturbation effects are largest for targets affecting growth and transcription and acting on highly expressed and variable downstream genes Our resource charts the impact of perturbing genes expressed or essential in iPSCs. Their knockdown resulted in a range of transcriptional change, with some targeted genes inducing hundreds of *trans* effects (**Figure 2A**). About half of the target genes (51%; 3,374/6,674) had at least one significant *trans* effect upon knockdown, with a mean of 2.57 per target gene. The genes with the highest number of significant *trans* effects have roles in a variety of fundamental biological functions including pluripotency maintenance, transcription and splicing. For example, members of the RNA polymerase associated factor (PAF) complex *PAF1*, *CTR9* and *RTF1*, had 962, 847 and 594 significant *trans* effects, while pluripotency maintenance complex members *CNOT1*, *CNOT3* and *myc*-associated factor *MAX* had 473, 281 and 144, respectively. These genes, which have previously been shown play important roles in the maintenance of pluripotency^19,20^, reflect the requirements for the stability of the core transcriptional circuitry in human iPSCs on the timescale of several days.

**Figure 2.**
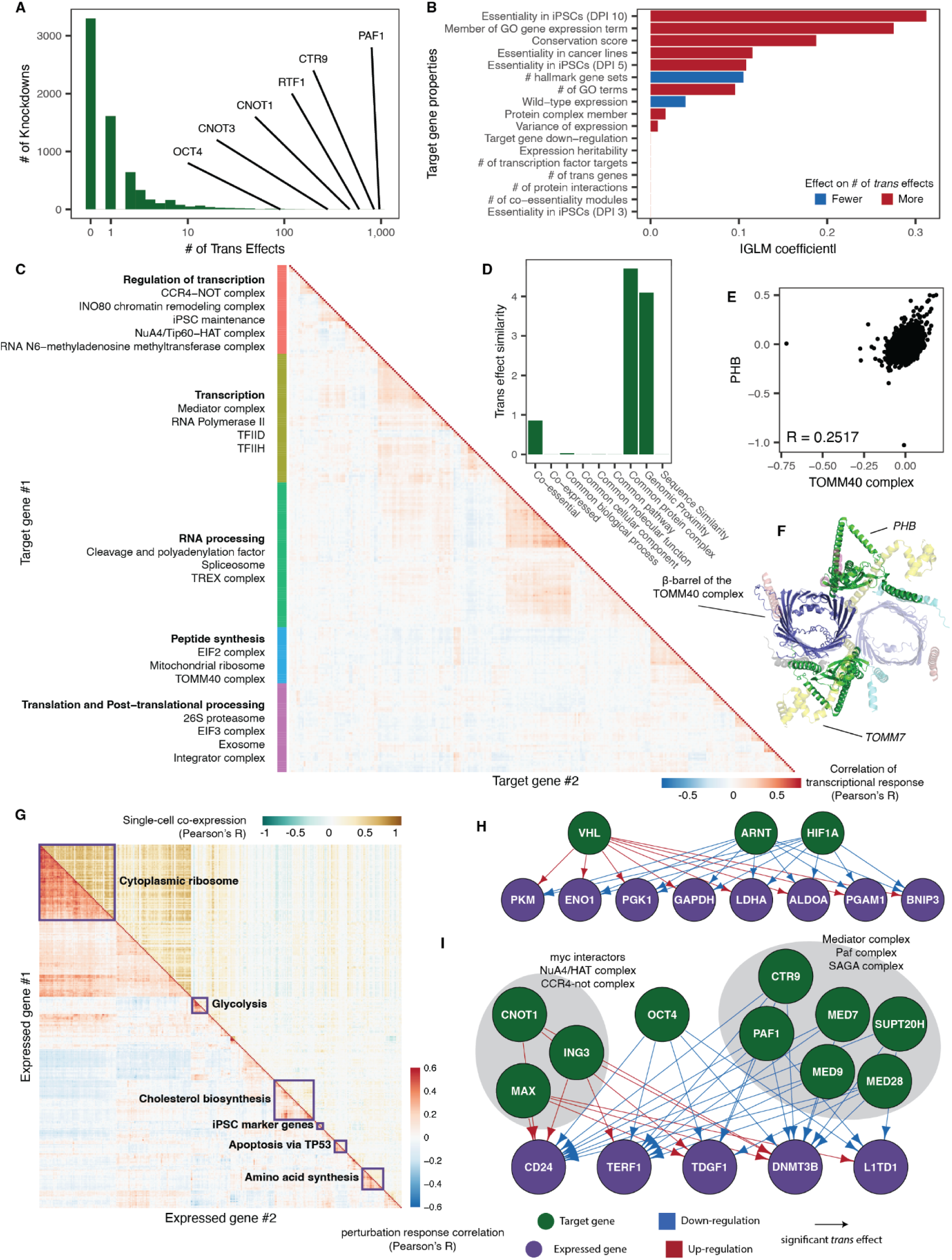
| Global transcriptional changes caused by gene-dosage reduction. A) Histogram of the number of significant trans effects per target. Target genes with the highest numbers are labelled. B) Model coefficients (x-axis) for predicting number of *trans* effects based on properties of the target gene (y-axis, DPI = days post infection). C) Heatmap of targets by *trans* effect similarity for the 280 knockdowns (with at least two co-regulating targets and at least one *trans* effect). D) Model coefficients (y-axis) for predicting similarity between *trans* effects based on functional relationship of two target genes (x-axis) E) Scatter plot of the *trans* effects induced by knockdown of genes in the TOMM40 complex (*x*-axis) and *PHB* (*y*-axis). F) Top view of an α-helix on *PHB* (green) predicted to interact with the β-barrel of the TOMM40 (blue) complex in a similar manner to known complex members, via a shared interface with *TOMM7*^37^ (yellow). G) Heatmap of correlation between expressed genes based on perturbation responses (lower-triangle) and natural single-cell co-expression (upper-triangle) for 313 expressed genes with the most co-regulated expressed genes (> 3 co-regulated genes). H) The regulatory behavior of hypoxia regulators *HIF1A*, *ARNT* and *VHL* on the glycolysis pathway. I) Regulators of commonly used iPSC marker genes. Knockdowns are indicated by green nodes while downstream genes are indicated in purple. Down-regulation upon knockdown is indicated in blue while up-regulation is indicated by a red arrow.

Next, we asked which properties of the targeted gene predict the number of significant *trans* effects they have. We included different biological features, as well as the number of assigned cells, in a regularized generalized linear model (**Methods**), identifying essentiality of the target as the most important feature, which is in line with findings in other cell types^21^ . This link was weaker for growth effects at earlier time-points (three and five days post-infection), as knockdowns with faster-acting growth defects could suffer from survival bias, with their fitness effects setting in before data were collected (**Figure 2B**). Among additional features, the number of Gene Ontology annotations, in particular being associated with gene expression^22^, as well as evolutionary conservation and protein complex membership of the target gene were also associated with a higher number of significant *trans* effects, an observation consistent with the hypothesis that constrained genes are required for diverse important functions^23,24^.

We then considered genes that responded in *trans* to the knockdown of one or multiple target genes (here-on referred to as regulators). Over half (67%, 4,365/6,471) of the expressed genes had at least one regulator, with an average of 2.6 (**Figure S2-1C**). Controlling for differences in expression level, the number of associated hallmark biological pathways^25^ and high expression heritability^1^ were the strongest predictors of having more regulators, while evolutionary conservation was the most informative predictor for having fewer (**Figure S2-1D**). These observations are consistent with previous hypotheses that genes with essential roles are more likely to be conserved across species and robust to perturbation of upstream regulators within one context, while genes with context-dependent expression levels are more responsive to regulation^23,24,26^ .

Next, we asked how the uncovered *trans* effects induced by CRISPRi knockdown could yield additional insights into the downstream regulatory consequences of natural genetic variants. Compared to identifying 512 significant *trans* effects by testing the expression of 2,044 genes across 26,877 SNPs in a collection of 1,367 iPSC lines^3^ (total of 230,010,344 tests; 2.2e-04%), our screen identified a total of 1,328 (3.5%) significant *trans* effects for the 5,127 genes with known *cis* eQTLs among the set of target genes. Out of the *trans* eQTLs identified previously, 67 pairs of *cis* and *trans* genes were also quantified in our data. However, we only found shared signal for one *trans-*eQTL, where the expression of *SLC3A2* was reduced upon knockdown of *SLC7A8*. The other *trans* eQTLs mapped in iPSCs did not yield a strong evidence of enrichment of significant expression change due to CRISPRi (**Figure S2-1E)**. The discrepancy between these two sets of trans effects could be explained by stronger effect sizes in the *cis* gene in our data (**Figure S2-2F**). Given the matching experimental cell line context, and the more severe expression perturbation in *cis* due to CRISPRi, this lack of replication of signal from a large natural population suggests that complementary assays such as CRISPRi or CRISPR activation can help to verify regulatory insights from existing *trans* eQTL studies and avoid false negative and positive signal.

Similarity of *trans* effects reveals protein complexes as the nexus for integrating gene expression changes Similarity of perturbation effect has proven to be one of the strongest lines of evidence for a functional link between two genes^9,27,28^. We therefore computed the correlation of gene expression changes upon knockdown for every pair of target genes across all expressed genes (**Figure 1E**). A rich tapestry of functional relationships emerged, that well recapitulated broad functional roles of the targets (**Figure 2C**). In addition to genomic proximity, which has been shown to be a strong predictor of off-target behavior^29,30^, protein complex co-membership was the strongest predictor of similar *trans* effects (**Figure 2D**). Indeed, knocking down individual members of a diverse set of protein complexes, such as the integrator complex, EIF3 complex and 26S proteasome, as well as complexes that have previously associated with the maintenance of pluripotency and self-renewal such as the eukaryotic gene expression regulator CCR4-NOT complex^31^ and histone modification complexes NuA4/Tip60-HAT^32^, RNA N6-methyladenosine methyltransferase^33^ and INO80^34^, induced similar expression changes (**Figure S2-2A-B**). Further, target pairs that were part of common co-essentiality modules defined by the Cancer Dependency Map^27^ also had more similar perturbation effects (**Figure 2C**). While these modules largely overlap with protein complexes, they additionally link complexes involved in similar functions. For example, knocking down members of RNA polymerase pre-initiation complexes TFIID and TFIIH had similar impact as knocking down members of the mediator complex and members of the RNA polymerase core complex itself (**Figure S2-2C**). This demonstrates how large-scale perturbation screening can link genes across hierarchies spanning from regulatory links to complexes and processes.

Consequently, we hypothesised that we could gain insight into the functions of poorly characterised genes by comparing their *trans* effects with those of better-studied protein complex members. We therefore aggregated cells with knockdowns from the same protein complex to obtain an average per-complex *trans* effect and computed its correlation to the *trans* effect of each target gene. We then prioritised 24 candidate gene-complex pairs based on correlation strength and shared known function or cellular compartment^35^, and predicted pairwise interaction structures between each target and all complex members using AlphaFold2-multimer^36^ to assess the biophysical plausibility of the interaction (**Supplementary Table ST2-1**). Complementary information from known complex crystal structures and other biological evidence can help determine if interactions are likely to occur.

Our candidate interactions form plausible complexes based on pDockQ^38^ scores significantly more frequently than random protein pairs (p < 10^-15^, Kolmogorow-Smirnow test) or known non-interacting pairs (p = 5 x 10^-13^), while following a similar distribution to known interactions from CORUM (p = 0.97)^39^. Eight of the 24 prioritized target-complex pairs had at least one plausible predicted interaction based on both pDockQ score and the visual inspection of the structures predicted by AlphaFold2, of which seven had a coherent interaction with the complex. This includes rediscovering the known structure for the interaction between *DDX39B* and *THOC2* in the TREX complex, and finding plausible interfaces in between *SMC3* and *MED16* from the mediator complex^40,41^ as well as *DDX41* and the tri-SNP complex^42–44^ via the homologous *LSM2*, *LSM5* and *LSM7* proteins^45^, cases where there is known to be an interaction but the structure of it is unknown. Beyond recovering known interactions, we identified four novel interactions with predicted plausible structures: *PHB* with the *TOMM40* complex (**Figure 2E-F**), *RAB10* with the Paf complex (**Figure S-2D-E**) and *ELP3* and *CTU1* with the EIF2B complex (**Figure S-2-2F-G**). These discoveries highlight the richness of functional data produced by genome-scale perturbation screens.

Genes with similar perturbation responses reflect activated cellular pathways and replicate naturally co-expressed gene clusters Similarity between expression changes of two genes in *trans* across different perturbations indicates shared regulation (**Figure 1F**). To analyse this effect in our data, we computed correlations between perturbation responses to all targeted genes for every pair of expressed genes and observed high values between members of various cellular stress pathways (**Figure 2G**). For example, downregulation of the master transcriptional regulator *HIF1A* and its binding partner *ARNT* caused down-expression of glycolysis genes *LDHA*, *GAPDH* and *ALDOA*, while knockdown of *HIF1A* degradation gene *VHL* resulted in the up-regulation of these genes, thus recovering the regulative relationship between hypoxia and the glycolysis pathways^46,47^ (**Figure 2H, S2-3A**). Similarly, targeting genes in the mevalonate arm of the cholesterol biosynthesis pathway^48^ resulted in the up-regulation of all other genes in the pathway (**Figure S2-3B-C**) indicating a shared feedback mechanism regulating their expression.

We next quantified which annotations best predict similarity of perturbation responses for all gene pairs and found co-expression of single-cell wild-type gene expression^49^ to be most informative (Pearson’s R = 0.41; **Figure S2-3D-E,** **Figure 2G**). In addition to cellular stress pathways and common protein complex membership (**Figure S2-3D**), which share signal with single-cell co-expression modules, this signal was driven by co-regulated expression modules, such as that linked to pluripotency maintenance (**Figure 2I**; **Figure S2-3F**). We observed co-regulation of a set of five commonly used iPSC marker genes, *CD24*, *TERF1*, *TDGF1*, *L1TD1* and *DNMT3B*^50–56^, which are also typically co-expressed in wild-type iPSCs^49^. The knockdowns that drove the shared signal were master pluripotency regulator *OCT4* and knockdowns with predicted off-target effects (**Supplementary Table ST2-2**, **Methods**) and components of the mediator and Paf complexes, which resulted in their down-regulation, and NuA4 complex member *ING3* gene, *CNOT1* and the *MAX* transcription factor, which resulted in the up-regulation of these genes, all of which have previously been attributed to the maintenance of stem cell pluripotency and development^20,32,57,58^. The finding of expression co-variation between individuals in natural populations to mirror common perturbation responses is of practical importance, as it allows designing efficient screening campaigns based on observational data. For example, savings on sequencing costs can be achieved by choosing a small set of genes for targeted capture, informed by correlation patterns in single cell co-expression, reasonably expecting to recover the main perturbation responses thanks to the shared signal.

### The influence of genetic background on gene perturbation effects

Gene perturbation effects can sometimes be suppressed or exacerbated by genetic background and small molecules^59^. We were therefore interested in understanding how often the *trans* regulatory effects that we observe differ in magnitude across healthy individuals, as this would indicate a mechanism for modulation.

To understand the malleability of gene perturbation impact by natural genetic variation, we capitalized on the diversity of donors of the induced pluripotent stem cell lines, and asked to what extent the variation of *trans* effects of a gene knockdown across cells could be attributed to genetic factors. We explored a targeted panel of 1,355 guides targeting 444 genes with 20 non-targeting guides, across 20 cell lines from 10 donors (2 lines per donor), focusing on genes more likely to have variable function across different cell lines (**Figure 3A**, **Methods, Supplementary Table 3-1**). After quality control, we recovered 1,161,864 cells of which we assigned 635,022 (55%) to 444 targets across 19 cell lines, collecting a median of 74 cells per target gene per line (**Figure S3-1A-B, Supplementary Table 3-2**). The knockdowns were again effective, with significant down-regulation of over 90% target genes in 15 of the 19 lines (**Figure S3-1C**). For every target gene, we computed *trans* effects on all 6,517 expressed genes (mean log-normalized expression > 0.1) across cell lines and for individual cell lines, following the analysis of the genome-scale screen (**Methods**). Across lines, the *trans* effects replicated the results from the genome-scale screen (Pearson’s R=0.73 across all *trans* effects that were significant in either screen), confirming that our earlier results were robust and reproducible (**Figure S3-1D**).

**Figure 3.**
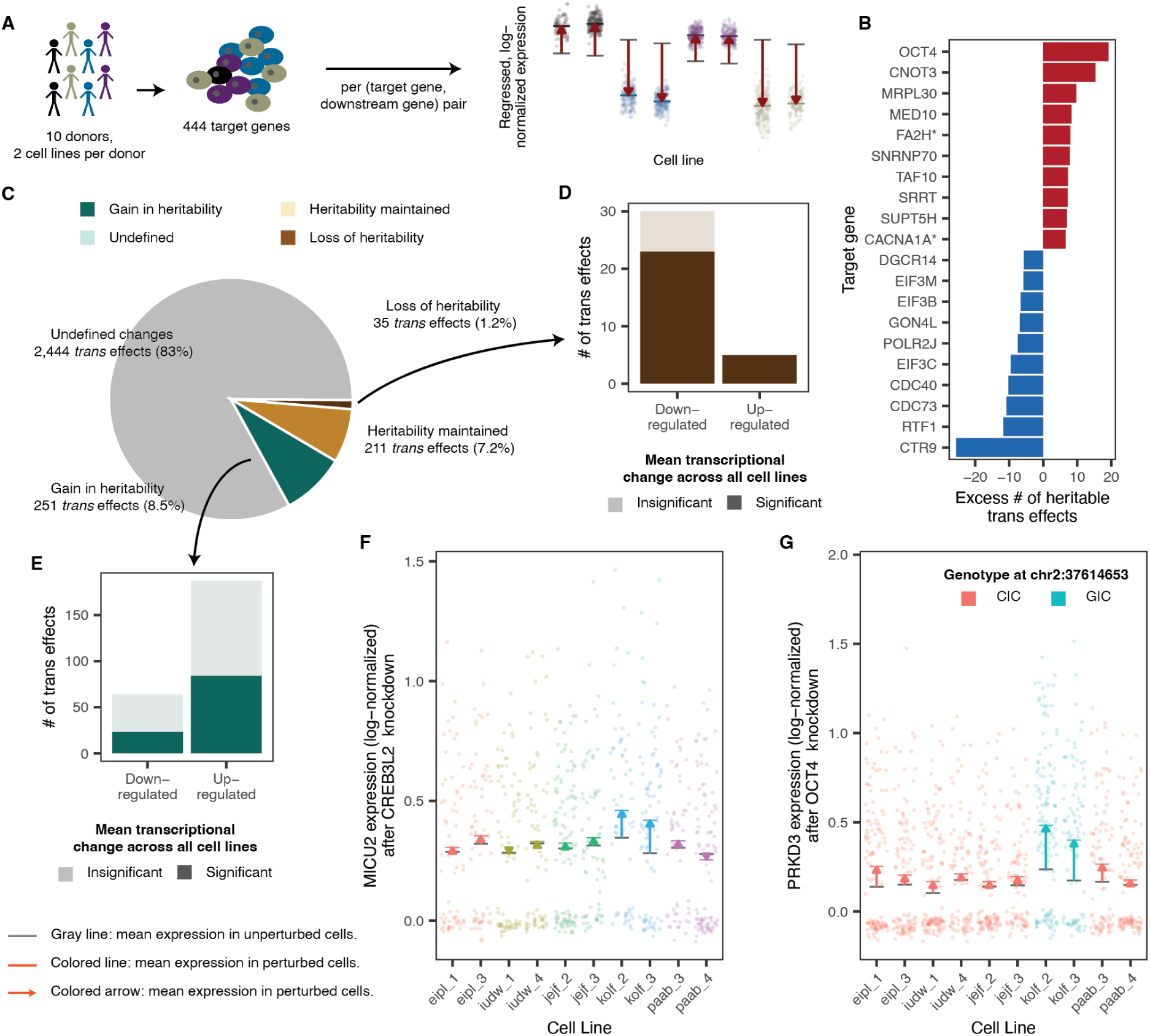
| Genetic background influences perturbation response. A) Analysis of variation in *trans* effects by considering a panel of 444 targets across 10 donors, with 2 cell lines per donor. B) Knockdowns with the most and least heritable *trans* effects, controlling for the number of *trans* effects tested. C) Breakdown of heritable, perturbation-induced, transcriptional change (out of 3,051). D) Mean transcriptional fold-change of *trans* effects resulting in loss of heritability of expression. E) Mean transcriptional fold-change of *trans* effects resulting in gain in heritability of expression. F) Expression change of *MICU2* due to knockdown of eQTL hotspot *CREB3L2*. G) Expression change of *PRKD3* due to knockdown of *OCT4*, separated by allele of a cis-eQTL of *PRKD3*.

When comparing perturbation effects in individual cell lines (**Methods**), we observed patterns consistent between the lines, with similar covariance structure between target genes (**Figure S3-2A-B**) and highly correlated response gene signatures (**Figure S3-2C**). To identify differences between genetic backgrounds, we considered only the 10 cell lines for the 5 donors where both cell lines from the donor indicated high response to CRISPR perturbation (**Figure S3-1C**) and focused on 68,321 *trans* effects (2.4% of all possible ones) that were significant in at least one cell line and could not be attributed to any off-target or cross-mapping effects (line-specific |LFC| > 0.1, Benjamini-Hochberg adjusted p-value < 0.1, **Methods**). We found 2,941 *trans* effects (4.3% of tested) for which the donor component explained a significant fraction of variance (Benjamini-Hochberg adjusted p-value < 0.1), which we call *heritable effects* (**Methods**). The knockdowns with the largest excess of heritable *trans* effects included pluripotency maintenance genes such as master pluripotency regulator *OCT4* and *CNOT3*; splicing genes *SNRNP70*, *SRSF1* and *ZMAT2*; as well as *FA2H* and *CACNA1A,* two genes whose misexpression has previously been attributed to the rare diseases hereditary spastic paraplegia and hereditary cerebellar ataxia, respectively^3^ . Conversely, *trans* effects due to knockdowns of PAF1 complex members *CTR9*, *RTF1* and *CDC73* and EIF3 complex members *EIF3C* and *EIF3B* were heritable less often than expected (**Figure 3B, Figure S3-3A**), indicating lack of segregating variation in the molecular impact of these key structures.

The interpretation of a heritable *trans* effect depends on the expression of the downstream gene in the unperturbed state (**Figure 3C**). If the gene expression was heritable in control cells (consistently variable across donors, Benjamini-Hochberg adjusted p-value < 0.1; 246/2,941 cases), heritable *trans* effects can emerge due to near-complete repression, or other outcomes that remove the variation between donors (“loss in heritability”, 35/246), such as for *C9orf135* expression changes due to *OCT4* knockdown (**Figure S3-3B**). Usually, this was marked by down-regulation of the expressed gene (28 out of 35, 80%, **Figure 3D**), though there were also several instances where a knockdown increased the expression of a gene in a single donor, removing expression differences between donors. An example for this is the up-regulation of mitochondrial cytochrome b (*MT-CYB)* in kolf_2 and kolf_3 cells upon knocking down various components of the large mitochondrial ribosome such as *MRPL55* (**Figure S3-3C**). Alternatively, expression heritability can be preserved where transcriptional change amplified effects of known genetic regulators like cis eQTLs or was small compared to the natural variation between donors (“maintenance of heritability”, 211/246) (**Figure S3-3D**).

In a second scenario, we considered heritability upon knockdown of the target genes to be potentially gained, if we could not find evidence for heritable expression in the control cells (2,695/2,941 cases, p-value > 0.9). Indeed, 251 of these cases showed significant evidence for heritable gene expression after the knockdown, and the majority (187/251; 75%) of them were a result of up-regulation in the expressed gene in a single donor (**Figure 3E**). Such *trans* effects would often be overlooked when testing across all lines where the dilution of signal results in insignificant *trans* effects for most of these cases (103/187, 55%). This gain in heritability can potentially be explained by genetic differences in the action of responsible genomic regulators, among other mechanistic reasons. We found that 3 of 7 previously mapped *trans* eQTL hotspot genes^3^ that we knocked down were linked to a heritable *trans* effect showing a gain in heritability in genes not previously linked to the eQTL: *CREB3L2* on *MICU2*, *ZNF208* on *SEH1L* and *ZNF611* on *MOB1A*, (**Figure 3F**; **S3-3F-G**).

Most heritable *trans* effects could not be directly linked to heritable expression before or after knockdown at the chosen significance levels. For example, the expression of *PRKD3* upon *OCT4* knockdown is heritable, and associated to a previously mapped *cis* eQTL at chr2:37614653 (**Figure 3G**). However, *PRKD3* expression heritability in our wild-type control cells could not be determined (p-value = 0.68) from the small number of donors. Larger screens with more donors will be required to categorize such heritable *trans* effects reliably.

### Towards design, implementation, and analysis of genome- and population-scale single cell CRISPR screens

The decrease of single cell library preparation and sequencing costs is making perturbation screening across a population, using a rich gene expression readout a reality. We have performed the first genome-scale screen with high-dimensional read-out across multiple individuals, and found that nearly all aspects of experimental design have an impact on the outcomes, which is important to consider in future studies.

In the targeted screen, guide identity had the largest effect on the observed *trans* effects among the genetic and experimental factors we tested (**Figure S4A-B**). While effects of different guides for the same target were mostly consistent (median Pearson’s R = 0.61 across all genes with at least 25 *trans* effects), unintended effects have been shown manifest from targeting similar sequences in other regions of the genome^60^, in particular where such regions lie in close proximity of a TSS of another gene^9,61^. Of 2,067 potential off-target *trans* effects, 112 exhibited significant repression of the predicted off-target gene and, on average, exhibited 24-fold greater variation due to guide than other *trans* effects (p-value = 1.3e-05, two-tailed t-test) (**Methods, Supplementary Table ST4-1**). Even stricter criteria might be needed to rule out all possible off-target effects. Perfect complementarity as short as 9bp of the guide seed sequence and promoter have been observed to result in off-target activity^61^ and such observations were replicated by those guides fulfilling this criteria on the expression levels of *OCT4* (**Figure S2-3F**).

In addition to guide identity, CRISPRi efficacy explained a large portion of variation in the observed *trans* effects (**Figure S4B**). The four lines where expression changes of the targeted genes that were not significant for over 50% of the library saw muted global transcriptional changes (**Figure S3-1C, S3-2C**). Both the CRISPRi on-target efficacy and the gRNA assignment rate were highly predicted by dCas9 expression in the cell line (**Figure S4C**), the latter likely due to the protective effect of the enzyme against degradation. These cell line effects indicate that successful screening requires a high dose of the Cas enzyme, and accounting for its efficacy in analysis, even after experimental selection for highly performing lines.

The experimental design required considerations on the number of cells per knockdown, approach to pooling cells from individuals, and timing of the assay were key to ensure that the resulting data were well-powered to draw reliable conclusions. First, depending on the sought effect size, anywhere from 10 to 200 cells may be needed to be profiled, as evaluated from a downsampling experiment (**Figure S1C**). Second, in an additional experiment knocking down 161 genes (**Supplementary Table ST4-2**) in 24 cell lines, and pooling prior to transfection of dCas9-KRAB-MeCP2, we observed that 6 out of 24 lines accounted for 98.4% of all the cells (**Figure S4D**), indicating technical challenge in ensuring balance across cell lines. Finally, in another experiment of 483 guides knocking down 161 genes with strong growth effects (**Supplementary Table ST4-2**) measured 14 days post-infection resulted in only 34% of cells successfully assigned to a guide, and no targets with significant downregulation (**Figure S4E**). In comparison, gRNAs could be successfully assigned for 52% of cells whose knockdown targeted a gene with weak growth effects, and 59% of the targeted genes had significant expression changes (**Figure S4E**). By sequencing earlier on days 3, 4, 5 or 6 post-infection and pooling cell lines late, we were able to recover sufficient numbers of cells in our later experiments (**Figure S1B**) and consistent target down-regulation (**Figure S2-1A**), highlighting the importance of carefully selecting the sequencing time-point when measuring impact of genes with fitness defects.

## Discussion

We have presented the first study in healthy humans to combine natural genetic variation with engineered perturbations and single cell sequencing readout. The impact of perturbations on gene expression ranges from limited impact to changing expression of thousands of downstream genes and highlights an overall concordance of observations from different research approaches, but also some differences. The main directions of variation in gene expression changes after knockdown are similar to those observed without perturbations in single iPS cells from a larger cohort^49^ (**Figure 2G**). Many of these co-expressed gene modules can be explained by a coordinated response to a signal, such as stress factors. When transcription factor perturbations had an effect in our system, they were always consistent with the annotated role of the factor as an activator or repressor. Lead *cis* eQTL SNPs explained substantial variation in heritable downstream perturbation effects as well. However, *trans* eQTLs, perhaps the most direct comparison to our study setup, mostly did not replicate, indicating either false positive mapping results for eQTLs, winner’s curse to diminish their effect sizes, false negative perturbation effects here, or discrepancies in experimental context. Cellular model resources with *trans* regulation maps as we have established here, power the analyses of *cis* eQTLs and expression-altering disease mutations by imputing their *trans* effects *in silico* and prioritising candidate genes.

Similar effects of perturbations indicated membership in the same protein complex or co-essentiality module. This allowed us to detect most complex members for several pieces of core cellular machinery. In addition, we could identify likely protein complex interaction partners by combining consistency of perturbation effect with AlphaFold2-enabled prediction that forecasts a confident interaction event. Similar screens in other cellular contexts where different complexes impact on gene expression could be used to identify their interaction partners or regulators.

Overall, protein complexes emerge as a nexus for integrating transcriptional changes and modulating downstream effects (**Figure 4**). Differences in gene expression, arising from factors like transcriptional stochasticity, cell cycle stages, epigenetic states, or genetic background, do not always lead to changes in protein abundance due to buffering mechanisms at the protein level^12^. For instance, excess uncomplexed proteins resulting from increased mRNA expression may be degraded due to exposure of otherwise hidden hydrophobic residues. Similarly, when rate-limiting components of protein complexes are degraded, the entire complex cannot form, leading to shared phenotypic impacts. These buffering mechanisms ensure that genetic effects on gene expression do not always manifest as changes at the protein level, thereby stabilizing cellular functions as reflected in the overall high degree of consistency in *trans* effects across cell lines. Together, our results indicate that protein complexes play a key role in mitigating the impact of gene expression variability, thus preserving the integrity of downstream processes.

**Figure 4.**
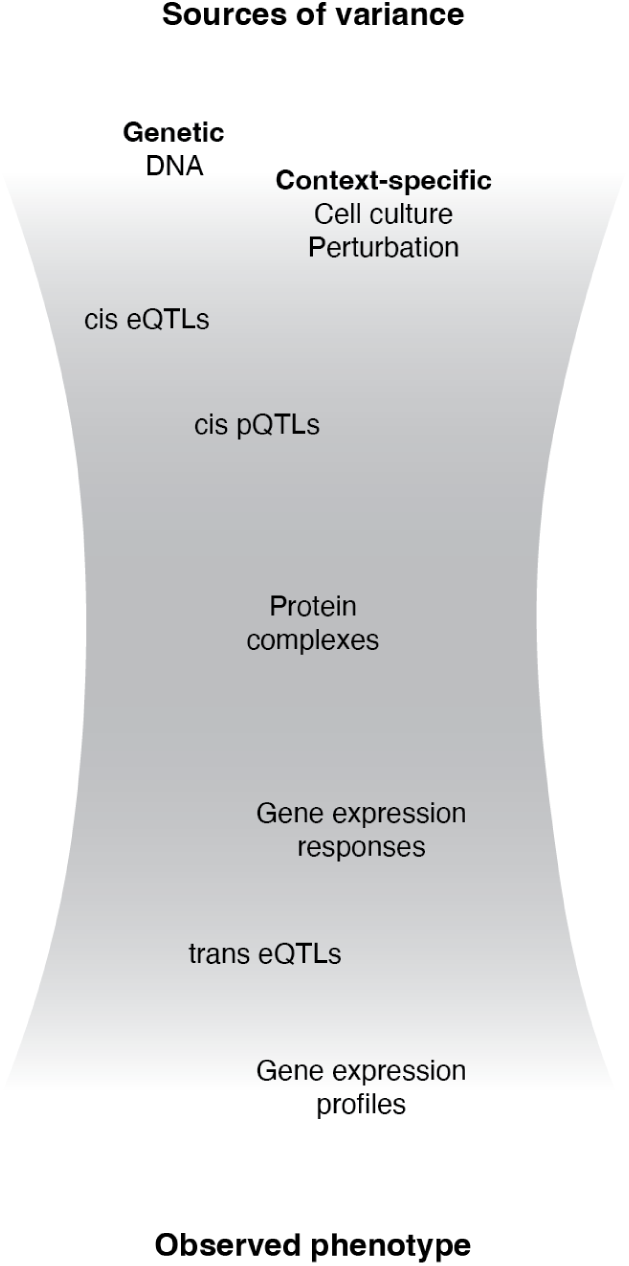
**| Robustness of knockdown effects to inter-individual differences.**

A venerable question in genetics is the extent of impact of modifiers, mutations that do not have large independent effects themselves, but modulate others. This is important both to be able to predict mutation impact, but also to identify the effects that can be modified at all, such that other ways to modulate cells, such as small molecules, could also be used. Many such alleles have been mapped in yeast, other model organisms, and human disease genes^62,63^. Our experimental design across multiple donors allowed us to identify perturbation effects that vary between individuals. Such heritable changes are small on average compared to technical impact of batches or sequencing coverage, as well as relative to biological covariates such as cell cycle state or CRISPR reagent efficacy. *Cis* eQTLs were frequently a likely natural cause for these effects, with the lead genetic variant associated to gene expression also influencing the response to perturbation.

The technological advances and cost reductions support scaling of genetic screening with single cell sequencing readouts. We have identified several factors that contribute substantially to the success of such campaigns, including strategy of pooling lines, sequencing time point, efficacy of engineering, reflected both by Cas9 activity, as well as on-target perturbation effect, gRNA recovery, and gRNA off-target effects. For CRISPRi specifically, off-targets are more promiscuous, with sequence matches in the 10nt of the PAM-proximal region sufficient to cause substantial unintended effects^61^. Large scale screening campaigns should map the impact of these variables separately before the start, in conditions that precisely match the scale-up phase later, to avoid complications.

Our study marks the beginning of genome-wide CRISPR screening with population-scale single cell RNA sequencing readout. This approach can be applied in cell models, organoids, and primary cells, where single screens have already provided new insights into human cell workings. Expanding these screens across many individuals and including more diverse genetic backgrounds will advance our understanding of disease causes and untangle genetic effects. While requiring careful technical optimization, such data are essential to building foundation models of individual-specific perturbation responses to genetic and small molecule changes, and ultimately predict and control cell behaviour.

## Acknowledgements

Grant support. C.F., E.M.P, Y.Z., K.C., L.C., S.U., A.D., J.S., M.E.S., D.M., E.G.N., A.B., S.L., Y.G., L.P., O.S., B.V. were supported by Wellcome (220540/Z/20/A). M.E.S was supported by Wellcome (220442/Z/20/Z). B.V., J.M.B were supported by the Deutsche Forschungsgemeinschaft (DFG, German Research Foundation) – 540147573.

## Author contributions

Designed project: L.P., C.F., B.V., L.C., Y.Z., E.M.P., S.U.

Performed experiments: E.M.P., Y.Z., K.A.X.C., S.C., A.B.

Analysed and interpreted data: C.F., B.V., A.D., M.J.B, J.M.B., J.S., M.E.S., D.M., E.G.N., S.L., Y.G., B.F., D.B.

Supervised study: L.P., O.S., B.V. Wrote paper: C.F., B.V., L.P., O.S., A.D.

## Resource availability

### Lead contact

Further information and requests for resources and reagents should be directed to and will be fulfilled by the lead contact, Leopold Parts.

## Materials availability

This study did not generate new unique reagents.

## Data and code availability

- Raw data for the genome-scale and targeted screen are available from SRA under the accession number ERP165335.
- Processed data, identified *trans* effects and heritability scores as well as additional data summaries to reproduce the findings from this manuscript are available from Figshare: https://figshare.com/s/14edeeab56eb8a885df3
- Code used for processing and analysis of the data is available from GitHub: https://github.com/claudiafeng123/crispri_scrnaseq_hipsci.
- An App for interactive exploration of the data is available here: https://www.sanger.ac.uk/tool/crispri-scrna-seq-hipsci/

## Supplemental information

● Supplementary Figures S1 - SM
● Supplementary Table ST1-1 : Selected genes for CRISPR knockdown in the genome-scale screen
● Supplementary Table ST1-2: Number of assigned cells per knockdown in the genome-scale screen
● Supplementary Table ST2-1: Plausibility of physical interactions between genes and complexes with similar transcriptional change due to knockdown
● Supplementary Table ST2-2: off-targets of *OCT4*
● Supplementary Table 3-1: Lines used for the targeted screen
● Supplementary Table 3-2: Selected genes for CRISPR knockdown for the targeted screen
● Supplementary Table 3-3: Heritable *trans* effects
● Supplementary Table ST4-1: Possible off-target effects based on less than 3 nucleotide mismatches to a Fantom 5 promoter
● Supplementary Table ST4-2: Genes selected for CRISPR knockdown in the pilot study
● Supplementary Table STM-1: Primers used
● Supplementary Table STM-2: Sequences used

## Methods

### Experimental design

#### Gene selection for the genome-scale panel

Guides were selected for the genome-scale panel based on wild-type expression levels in iPSCs, as well as those that demonstrated growth effects. Guides were then separated into those with targeting genes with fitness effects (fitness genes), which were measured at 3, 4 and 5 DPI, and those without (non-fitness genes), which were measured at 6 DPI. 40 non-targeting control guides from Dolcetto A were used for both panels.

To identify genes with fitness effects, we conducted an essentiality screen using the Dolcetto library^17^. We used 57,050 guides targeting 18,899 genes from the Dolcetto A library and 57,011 guides targeting 18,897 genes from the Dolcetto B library to knock down a total of 18,940 genes in *fiaj_1* cells and harvested cells at 3, 4, 5, 9 and 10 days post-infection. At each time-point, fitness effects were quantified by calculating the log_2_-fold change of normalised cell counts compared to that of the read counts in the plasmid library^64^ and genes were considered to have fitness effects if the median fitness effect at day 10 across all guides was less than -1. The three guides with the lowest log_2_ fold change at day 10 post-transfection were then chosen for screening. If fewer than 3 guides were available across both Dolcetto A and B libraries, all available guides were chosen. In total, this part of our library consisted of 6,784 guides targeting 2,264 targeted fitness genes.

Additional genes, without a fitness effect iPSCs were selected based on fitness effects in cancer cell lines, as well expression level in iPSCs. Fitness effects in cancer cell lines were assessed based on the CERES scores of all 1,376 lines in the DepMap consortium^16^. Gene expression in iPSCs was measured in a pilot screen on *fiaj_1* cells and expression values per cell were normalised by total sum scaling with a scale factor of 100,000 and log-transformation with a pseudo-count of 1^65^. Additional targets were considered if they were either highly expressed in iPSCs (normalised expression > 0.1), or had strong fitness effects in cancer cell lines and were expressed in iPSCs (genes with a CERES score > 0.22 or <-0.25 on average across all lines or in at least 100 lines and and a normalised expression > 0.01) or had variable fitness effects (genes with a CERES score standard deviation across lines > 0.15 and a normalised expression > 0.01). In addition, the 50 genes with highest CERES score standard deviation but expression < 0.01 were selected. For each gene, 3 guides were selected from the Dolcetto A library, complementing with guides from the Dolcetto B library if less than three guides were available in Dolcetto A. In total, 4,962 genes and 14,883 guides were selected.

#### Gene selection for the targeted panel

Genes were selected for the targeted panel for measuring genetic background effects based on their effect size observed in the genome-scale screens. Only target genes with at least 20 cells per gene, log-normalized expression greater than 0.1 and minimum correlation of significant *trans* effects across timepoints and guides larger than 0.5 were considered for selection. Of these, all genes with more than 12 significant *trans* effects were selected for the panel (n=110 genes, 106 of which were fitness genes). For comparison, we added 97 genes with 5-12 significant *trans* effects (all of which were fitness genes) and 203 genes with fewer than 5 significant *trans* effects (141 of which were fitness genes). In addition, we considered 7 genes that have been linked to eQTLs with many *trans* effects^3^, 38 genes associated with monogenic diseases^3^, 17 genes with high variance of CERES scores across lines^16^, 15 genes with a high expression heritability^1^ and *ARID1A*, *EZH2* and *BCOR*. Guides were chosen as in the genome-scale screen apart from 5 outlier guides, for which the target gene log-fold change was higher than the upper bound of a 95% confidence interval in a linear regression of guide-level versus target-level log-fold changes and a suitable replacement guide with a similar growth-effect ten days after transfection could be found in the Dolcetto libraries. This resulted in a total of 1,355 guides targeting 444 genes and 20 non-targeting control guides for the targeted panel.

### Experimental protocol

#### Molecular cloning

The libraries were cloned into the lentiviral expression library pKLV2-U6gRNA5(BbsI)-ccdb-PGKpuroBFP-W (Addgene 67974)^66^. Briefly, the guide libraries were ordered from Twist Biosciences as 215-mer oligo pool. The pool was composed of several sub-pools to allow for the selective amplification of gRNAs that were amplified with subpool specific primers (**Supplementary Table STM-1**). BBSI-digested amplicons encoding gRNAs were inserted into the BBSI-digested vector by Gibson assembly (NEB Gibson Assembly Master Mix) according to manufacturer’s specifications, and transformed by electroporation (NEB 10-beta Electrocompetent E. coli C3020K). Bacterial cells were cultured overnight in liquid culture and plasmid DNA was extracted. The plasmid libraries were pooled together in equimolar ratios to achieve the desired final libraries.

For the construction of the pB-CAGGS-dCas9-KRAB-MeCP2-BSD-mScarlet plasmid, the pB-CAGGS-dCas9-KRAB-MeCP2 (Addgene 110824) vector was digested with NotI (NEB) and EcorV (NEB). The EF1α promoter and blasticidin resistance gene was amplified by PCR using primers #1009 and #1010 (**Supplementary Table STM-1**). The SNV40 polyA signal was amplified by PCR using primers #1013 and #1014 (**Supplementary Table STM-1**). The mScarlet sequence was amplified by PCR from plasmid pmScarlet_C1 (Addgene 85042) using primers #1016 and #1012 (**Supplementary Table STM-1**). All products were purified with Monarch DNA Cleanup Columns (NEB). T2A sequence was ordered as a gBlock from IDT. A Gibson assembly with 4 fragments is incubated at 50°C for 30 minutes and transformed by electroporation.

#### Cell culture

Human iPSCs were cultured on Vitronectin XF (StemCell Technologies, 07180)-coated plates and mTeSR Plus medium (StemCell Technologies). The medium was changed every other day throughout expansion and all experiments. Cell lines were cultured at 37°C, 5% CO_2_.

#### dCas+ cell line generation and activity validation

For the generation of dCas9-KRAB-MeCP2 iPS cell lines, *3 x 10*^5^ wild type cells were seeded into 12-well plates with ROCKi containing media. For the transfection of one line, 600ng of pB-CAGGS-dCas9-KRAB-MeCP2-BSD-mScarlet, 300 ng of mPBase^67^ and 100ng of a reporter plasmid encoding for GFP were mixed with 50ul of Opti-Mem in one tube and 50ul of Opti-Mem was mixed with 2 ul of Lipofectamine Stem (Invitrogen) in another tube. After 5 minutes of incubation at room temperature, the contents of the tubes were mixed together and incubated for another 10-30 minutes at room temperature. During incubation, the media in the wells was refreshed and 0.5ml media was added. After incubation, 100ul of the complexes were added to the wells. 24h after transfection, 1ml of media was added to cells. 48h after transfection, blasticidin (TOKU-E) selection was started using a concentration of 2µg/ml. The cells were cultured in selection for 2 weeks.

To validate the dCas9-KRAB-MeCP2 activity of the cells, an adopted method of the previously published Cas9 validation system was used^64^. Briefly, cells were transfected with a plasmid that encodes for BFP and GFP and either a mock gRNA or a gRNA targeting GFP TSS. *1 x 10^5^* cells were seeded into 24-well plates. Cells were transfected with either the mock or silencing construct using Lipofectamine Stem 24h later. BFP and GFP expression were measured three days after transfection at FACS. dCas9-KRAB-MeCP2 activity was calculated based on the median expression of GFP in BFP positive cells. Two replicate measurements were made for all cell lines for both conditions.

#### Lentivirus production and determination of lentiviral titer

Supernatants containing lentiviral particles were produced by transient transfection of 293FT cells using Lipofectamine LTX (Invitrogen). 5.4 μg of a lentiviral vector, 5.4 μg of psPax2 (Addgene 12260), 1.2 μg of pMD2.G (Addgene 12259) and 12 μl of PLUS reagent were added to 3 ml of OPTI-MEM and incubated for 5 min at room temperature. 36 μl of the LTX reagent was then added to this mixture and further incubated for 30 min at room temperature. The transfection complex was added to 80%-confluent 293FT cells in a 10cm dish containing 10 ml of culture medium. After 48 h viral supernatant was harvested and fresh medium was added. After 24h the lentiviral supernatant was collected and mixed with the first supernatant which was then stored at -80°C.

For gRNA library lentiviral titration on dCas9-KRAB-MeCP2 expressing iPSCs, iPSCs were harvested by Accutase (Stemcell Technologies) as single cells. iPSCs (3.6x10^5^/well in 6-well plate) were infected with at least five serial dilutions of lentiviral supernatant supplemented with 10µM Rock inhibitor Y-27632 (Stemcell Technologies). Uninfected cells were used as negative control. The transduced cell mixture was cultured in 6-well plates in 2ml/well. 24h post transduction, the medium was refreshed with mTeSR Plus without Rock inhibitor. After three days of cell culture the cells were harvested for FACS analysis and the level of BFP expression was measured. Virus titer was estimated and scaled up accordingly for subsequent screens.

#### Screening and sequencing

Cells were transduced with the lentivirus aiming for an MOI of 0.2. The cells were seeded at a density of *2.0 x 10^5^* to *4.5 x 10^5^* depending on the day of harvest. Media was refreshed 24h after transduction. Cells were harvested either on day 3, 4, 5 or 6 after transduction. On collection day, cells were harvested with accutase, spun down and resuspended in eBioscience Fixable Viability Dye eFluor 780 (Invitrogen) that was diluted 5000-fold in eBioscience™ Flow Cytometry Staining Buffer (Invitrogen). Cells were stained for at least 5 minutes and then filtered with Scienceware® Flowmi™ Cell Strainer (SP Belart). Cells were then sorted based on dead/alive-staining, BFP and mScarlet expression on MA900 Multi-Application Cell Sorter (Sony), The BD Influx™ (BD Biosciences) or MoFlo XDP Cell Sorter (Beckman Coulter). An equal number of cells were sorted for all the lines. 12 lines and 8 lines were pooled together for the genes with and without fitness effects, respectively, and *1.65 x 10^4^* cells were loaded in a 10X inlet. Chromium Next GEM Single Cell 5’ Kit v2 (10X Genomics) was used for transcriptome capture, with a modified protocol where we added an extra primer to the GEM generation mix to capture gRNAs^21^.

### Computational analysis

Unless otherwise stated, all analyses were performed in R (version 4.3.1)^68^ and Seurat (version 5.0.3).

#### Read alignment using CellRanger

Reads were aligned with CellRanger^69^ (version 6.0.1), processing each inlet separately. Alignment was conducted using default parameters, using genome build GRch38 as a reference, an adding additional sequences for BFP, mScarlet, BSD and dCas9-KRAB-MeCP2 (**Supplementary Table STM-2**). The sgRNAs were aligned to libraries for the fitness genes and other genes, respectively. For one inlet, the minimum threshold for the GEX/Cite-Seq cell barcode overlap was lowered from 0.1 to 0.01.

#### De-multiplexing of cells based on natural genetic variation

Individual cells were assigned to the source cell lines by de-multiplexing using natural sequence variants, as each pool consisted of lines from different individuals. We first used cellSNP 0.1.7^70^ to call genotypes from the bam files containing the 10x read sequences for all cells passing the CellRanger filters. We used bcftools^71^ (version 1.10.2) to subset a list of candidate SNPs^72^ to only lines present in each inlet and filtered for a min. allele frequency threshold of 0.01 and minimum aggregated count of 20. This output was used in Vireo (version 0.2.1)^73^ to de-multiplex the cells into the number of lines present in each pool using genotype data for each donor provided by the HipSci consortium^1^, modified variant coordinates from GRCh37 to the genome build GRch38 using CrossMap^74^. In total, 69% and 72% of cells were uniquely assigned to one cell line in the genome-wide and targeted screens, respectively. Doublets and unassigned cells were removed for further analysis.

#### Quality control and filtering

High quality cells were retained based on three criteria: number of RNA UMI counts per cell, number of unique features per cell and percentage of mitochondrial RNA^75^. The number RNA UMI counts and unique features per cell were either bimodal or trimodal for each inlet and we removed cells that were in the lowest mode of number of features and UMI counts using inlet-specific thresholds between 1,926 and 39,260 UMIs per cell (average of 14,186 RNA UMIs across inlets), 2,000 features per cell and a percentage of mitochondrial genes above 10%. After filtering, we assigned cell cycle scores for each cell using Seurat’s *CellCycleScoring* function with cell cycle marker genes retrieved as *cc.genes.updated.2010*^65^.

#### Guide assignment

To establish an optimal guide assignment strategy, known to impact power and discoveries^76^, we considered a pilot data-set knocking down **161** genes with weak fitness effects in 24 iPSC lines (**Supplementary Table ST4-1**). We employed five different tools and evaluated the quality of each assignment by considering the number of knockdowns with significant on-target down-regulation and the median number of cells per guide (**Figure SMA-B**). Based on this, we considered the relative UMI abundance of the most abundant guide with respect to the total number of guide UMIs in a cell for guide assignment in all further analyses. Cells were assigned to a guide if the relative frequency of the most abundant guide was in the upper mode across cells within a cutoff window 0.5 and 1 (median threshold across inlets was 0.75, minimum 0.5 and maximum 0.88) and had a minimum of 3 UMIs in the cell. All other cells were considered unassigned. The percent assigned cells varied across inlets and library sizes, ranging from 9% to 78% in the genome-wide experiments and 37% to 69% in the targeted experiments, with an average of 30% and 55% of cells assigned to a single guide, respectively.

#### Data integration and variance component analysis

For all cells passing quality control we normalised the data by total sum scaling with a scale factor of 10,000 and log-transformation with a pseudo-count of 1 and combined these results using Seurat’s *merge* function, keeping all genes with a minimum normalised expression of 0.1 in all inlets, resulting in a total of 6,471 expressed genes in the genome-wide screen and 6,517 expressed genes in the targeted screen. To produce UMAP plots, we extracted highly variable features using *FindVariableFeatures*, performed PCA using *RunPCA* and calculated a UMAP embedding on the top 20 PCs. To quantify the contribution of the different variables on the transcriptome heterogeneity, we used a linear mixed model on the expression of the 2,000 most highly variable genes in a variance component analysis including donor / cell line, batch / inlet, cell cycle phase, sequencing time point and target gene as random effects and percentage of mitochondrial genes and total number of UMIs per cell as fixed effects. To remove technical and batch differences as well as line-specific effects, the corresponding variables were regressed out from the normalised expression data using *ScaleData* with *vars.to.regress* set to the respective variables and the PCA and UMAP were re-calculated on the residuals of the model.

#### Quantifying perturbation effects

To quantify perturbation effects in the genome-wide screen, we defined all unassigned cells as well as cells assigned to a non-targeting guide as control cells. Based on the gene expression measurements of all control cells, we used a linear model to estimate the effects of cell line, inlet, percent of mitochondrial genes, cell cycle scores and total number of UMIs per cell on the gene expression. To assess the effect of a perturbation within an assigned cell, we calculated the expected expression of each gene based on the linear model and compared this to the observed expression, yielding a *perturbation effect profile* for each cell defined as the difference of the expected and observed expression.

To assess overall perturbation effects per guide, per target or per target x line pairing, we averaged these effects across all cells assigned to a guide, target or target x line pairing, respectively, with significance evaluated based on a *z*-test using the residuals variance of the control fit. For the genome-wide screen, targets were considered for analysis if they had a minimum of 10 cells assigned to it, individual guides if they had a minimum of 5 cells. This left a total of 14,982 guides and 6,673 targets to be considered, for which *trans* effects on 6,471 genes were calculated, giving a total of 96,948,522 and 43,180,983 *trans* effects across guides and targets, respectively. For the targeted screen, effects for each of the 444 knockdowns were computed across all lines, as well as separately for each of the lines with where there were least 10 assigned cells (total of 8,204 *trans* effects). As in the genome-wide screen, only the transcriptomic changes for the genes whose log-normalized mean expression was greater than 0.1 were computed (6,517 expressed genes). In total, effects were computed across 53,465,468 target, line, expressed gene triplets.

For *cis* effects by natural genetic variants, we used the same procedure to estimate *cis* effects size on the known *cis* gene,using all control cells in the model and replacing the cell line covariate with the number of alternate alleles as a proxy for the genotype of each donor. Cis effect sizes for every eQTL were determined as the model coefficient.

*Trans* effects were considered significant if the *p*-value after Benjamini-Hochberg correction across all targets, tested genes and lines (if applicable) was below 0.1.

#### Power estimation based on down-sampling experiments

We estimated the variance of the estimated transcriptional change due to knockdown and impact of the number of assigned cells using a bootstrap procedure. For this, we considered 118 targets of varying effect sizes (11 < # of differentially expressed genes < 551) where the full data set had at least 1,000 assigned cells. For each target, we subsampled all cells with replacement to obtain a simulated dataset of 5, 10, 25, 50, 100, 250, 500 and 1000 cells. Transcriptomic changes for all expressed genes were then computed separately on each of these data sets exactly as on the full data set (see *Quantifying perturbation effects*) This was done 25 times per knockdown, resulting in a total of 25 separate estimates of transcriptional effect for 8 different sample sizes for each of the 118 targets.

#### Identifying co-regulated and co-regulating genes

To quantify the similarity of targets (and expressed genes) based on their perturbation effects (perturbation response) we calculated Pearson’s correlation between targets (expressed genes) based on the log-fold changes for all 6,471 expressed genes (6,673 well-powered targets). A total of 22,261,128 target-target and 20,933,685 expressed gene-expressed gene pairs were considered for analysis. Two targets (expressed genes) were considered to be co-regulating (co-regulated) if the absolute Pearson’s correlation between their *trans* perturbation (response) was greater than 0.2.

#### Quantifying heritability of perturbation effects across donors

Heritability was estimated in the targeted screen, considering the 5 donors where both cell lines from each donor demonstrated strong response CRISPR perturbation. We considered every target-expressed gene pair where we observed a significant *trans* effect in at least one line (68,321 target x expressed gene pairs). For each of these pairs, the *perturbation effect profile* for every assigned cell obtained from *Quantifying Perturbation Effects* was fitted with a linear mixed model using normalized target gene expression as a fixed effect and guide, cell line and donor as random effects. Donor effect was quantified by computing the likelihood ratio between the full model and the model without donor as a random effect. To assess significance we created a permutation scheme to obtain an empirical null distribution of the donor effects. For this, donor labels were permuted across cell lines such that two cell lines from one donor were assigned to different donors after the permutation. Thereby we retain the cell line structure in the data but permute the donor structure. This yielded a total of 544 permutations for the 5 pairs of cell lines. Permutations for every target-gene pair were computed until 10 null values greater than or equal to the true value were observed (*stronger observations*), or until 10^4^ null values were computed. The empirical p-value was estimated to be *p_td_ = (*max(10, # Stronger Observations *+ 1)/(*min(# of permutations, 10^4^) *+ 1)*. Effects where the Benjamini-Hochberg adjusted p-value < 0.1 were considered to be significant.

All models were fitted using the lmer function from the R package lme4 (v1.1-35.1)^77^. Log-likelihood was computed using the logLik function from the R stats package.

#### Identification of off-target effects

A knockdown was considered an off-target effect based on two criteria:

- **Genomic proximity**: guide sequence could be mapped to within 1kbp of a transcription start site of another gene in the Fantom 5 database^78,79^
- **Sequence similarity**: any sequence with fewer than 3nt mismatches could be mapped to within 1kbp of a transcription start site of another gene in the Fantom 5 database^78,79^.

To identify guides with potential off-target effects on *OCT4*, we additionally considered any guide in our library whose first 9nt of their seed sequence could be mapped to a region within 2kbp of a transcription start site of *OCT4* in the Fantom 5 database.

#### Functional annotations

Functional annotations were used throughout analysis, such as for predicting number of *trans* effects and number of regulating target genes, similarity of transcriptional effects upon target downregulation and co-perturbation. To do this, we made use of the following annotations:

### Conservation scores

Conservation scores were obtained for each target from the Bioconductor package phastCons100way.UCSC.hg38 (version 3.7.1)^23^.

### Wildtype expression, correlation and variance of expression

Wild-type gene expression, variance and co-expression was calculated based on undifferentiated iPSCs^49^. Values were computed based on the log-normalised expression values after regressing out effects due to donor and technical covariates (percent of mitochondrial genes, total number of UMIs per cell and number of genes expressed). Two genes were considered *co-expressed on the single-cell level* if their absolute Pearson correlation was above 0.3.

### Essentiality

Essentiality was quantified as described in *Gene selection for the genome-scale panel* based on an iPSC cell line and the DepMap consortium^16^.

### Expression heritability

Heritability of wild-type expression of iPSCs was obtained from the HipSci consortium^1^.

### Protein-protein interactions

Known gene interactions were obtained from the OmniPath database^80^ using the *import_all_interactions* function from OmnipathR (version 3.10.1)^81^. A pair of genes was considered to be interacting if they formed an interaction pair in the OmniPath database (undirected).

### Protein complexes

Known protein complexes were obtained from the Omnipath database^80^ using the *import_omnipath_complexes(resources = c(’CORUM’, ’hu.MAP’))* function from OmnipathR (version 3.10.1)^81^. Pairs of genes were considered to be a protein complex pair if they were both members of at least one common protein complex.

### Transcription factor regulation

Transcription factor-target gene interactions were obtained from the DoRothEA database^82^ using the function *get_dorothea* in the decoupleR package (version 2.8.0)^83^, using all pairs with confidence level A or B.

### Hallmark gene sets

Knockdown and target genes were annotated by their membership in release 7.5.1 of the mSigDB hallmark gene sets^25^. Two genes were considered to be a hallmark gene set pair if they were both members of at least one gene set.

### Co-essentiality

Co-essentiality modules were taken from Wainberg et. al.^27^. Two genes were considered to be a co-essentiality module pair if they were both members of at least one co-essentiality module.

### GO Terms

GO term annotations were obtained from the authors of the gProfiler database^84^. Pairs of genes were considered to be an enriched gProfiler pair if they had at least 1 GO annotation in common.

### Expression Quantitative Loci in Human iPSCs

Putative *trans-*eQTLs were obtained from^3^ on February 27, 2023. A pair was considered a *cis* eQTL*-trans* eQTL pair if the target gene and identified *trans* gene in our data corresponded to a *cis* eQTL gene and its *trans* effect.

Unless otherwise stated, prediction was done by fitting an elastic net regression model with *alpha = 0.1* using of the *glmnet* package (version 4.1-8)^85^. When predicting the discrete, positive values (i.e. the number of *trans* effects and regulating knockdowns), a Poisson regression was used. When predicting binary response (i.e. similarity of perturbation response or perturbation profiles), a logistic regression was used. To account for differences in statistical power we controlled for the number of cells by adding this as a term in the model.

#### Protein Complex Prediction

We identified candidate complex interactions based on the correlation between candidate target genes downstream effects and those of known complex members (Pearson’s R > 0.2). Candidates were then categorised into known and novel interactions based on literature review and prioritised according to shared function and cellular compartment with the complex. AlphaFold-Multimer^86^ (version 2.3) was used to model pairwise interactions between each target gene and all members of the candidate complex. pDockQ^38,87^ scores were calculated for each interaction as well as a random background sample of protein pairs and a set of known protein interactions. We identified plausible target-complex interactions with a combination of manual examination of predicted structures and pDockQ scores. We then aligned the top predicted pairwise target-complex member interactions with known complex structures using PyMol^88^ (version 2.5 Open-Source).

#### MOFA

Multi-modal factor analysis was used to compare *trans* effects across cell lines using MOFA2 (version 1.12.1)^89,90^. For this, log-fold change values for 445 target genes, expression of 6,517 genes, dCas9-KRAB-MeCP2, BSD and mScarlet and 19 cell lines were z-transformed and input to MOFA with default parameters, with each cell line as a separate view in the model and the number of factors set to 8.

---

**Figure S1.**
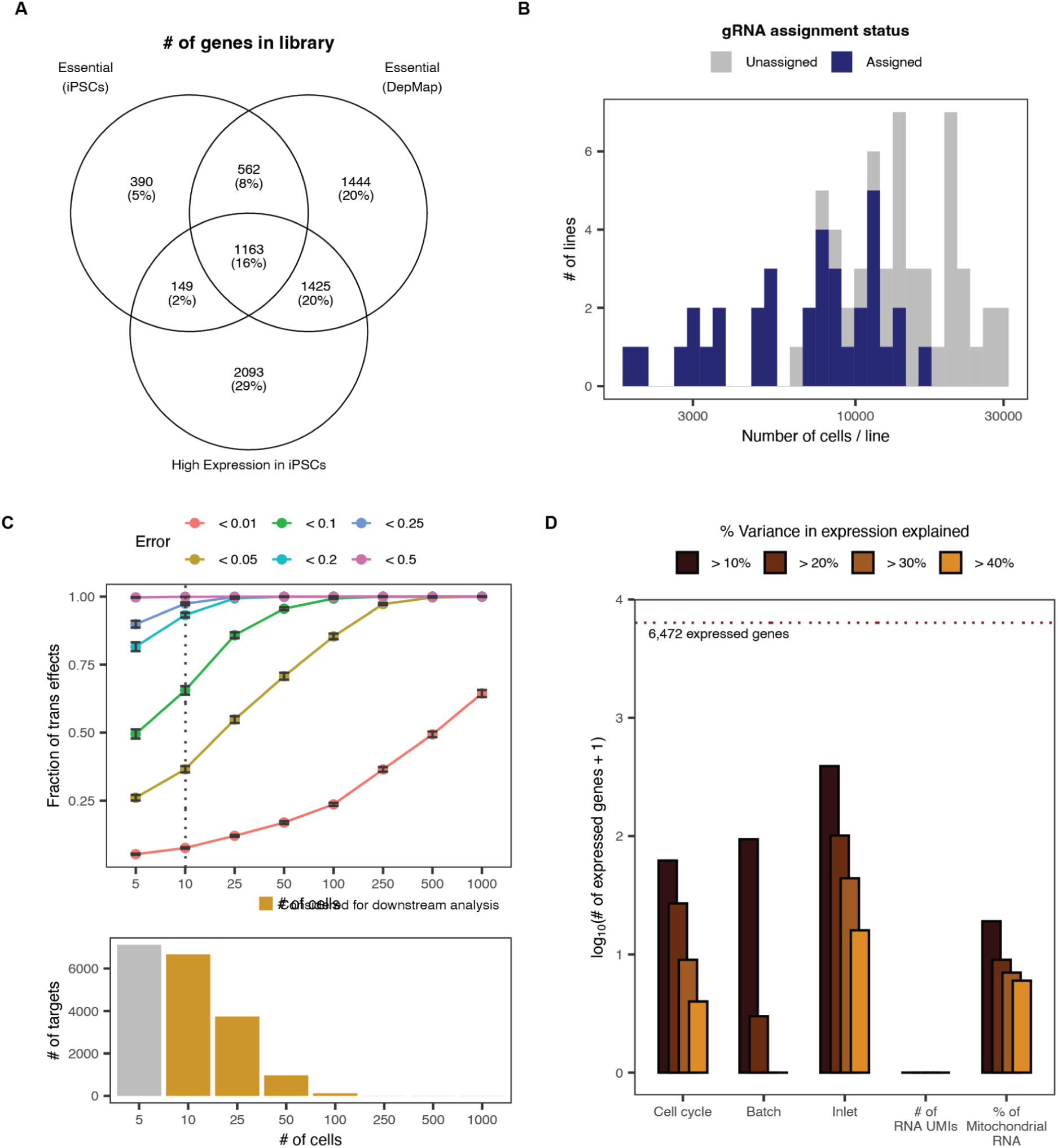
| Coverage and variance in the genome-scale screen. A) Venn diagram of genes selected as targets. B) Number of lines (y-axis) for different numbers of cells recovered per line after genotyping (x-axis) with gRNA assigned (blue) and not (grey). C) Estimated absolute error of expression log-fold changes for varying number of assigned cells (top) (relative to estimates from all genes, with a minimum of 1,000; Methods), histogram of cells per target gene (bottom). D) Number of genes (y-axis; log10 scale) with increasing amounts of variance explained (colors) by cell cycle and technical artefacts (x-axis).

**Figure S2-1.**
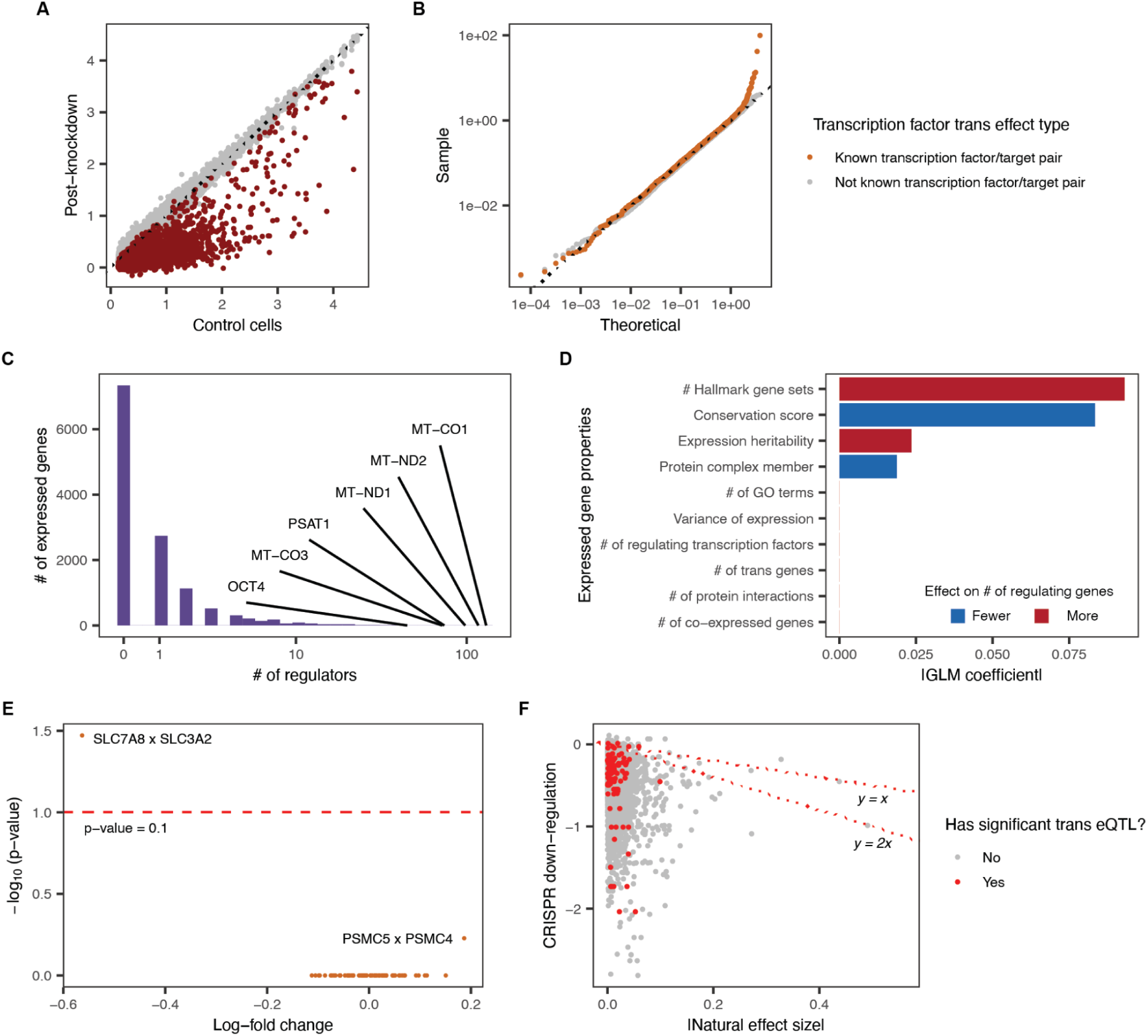
| Molecular signatures of the *trans* effects of gene knockdown. A) Down-regulation of target genes due to CRISPRi. Expression of targeted (red markers) and other genes (grey markers) Red dots show target gene expression values in control cells (x-axis) and assigned cells (y-axis), grey dots show expression values of expressed genes beyond the target. B) Quantile-quantile plot of *p-*values of *trans* effect of transcription factors in the DoRoTHeA database. Orange points indicate the *trans* effects of transcription factors and their known targets while gray points indicate *trans* effects the same transcription factors with non-targets. Number of downstream genes (y-axis) with different numbers of regulators (x-axis). Labels: six genes with most upstream regulators. C) Histogram of the number of regulators per expressed gene. Expressed genes with the highest numbers of regulators are labelled. D) Absolute model coefficients (x-axis) for predicting the number of regulators based on properties of the expressed gene (y-axis). Blue: negative coefficients (fewer regulators); red: positive coefficients (more regulators). E) Volcano plot of the log-fold change (x-axis) and log-scale significances (Benjamini Hochberg adjusted p-values, y-axis) for *trans* effects of (target, expressed gene) pairs with a known eQTLs acting in *cis* on the target and *trans* on the expressed gene. Dashed line: p=0.1. Labels: two pairs with corrected p-value less than 1. F) Comparison of effect size of natural variation in expression attributed to a cis eQTLs and CRISPRi. Red: cis eQTLs with at least one significant *trans* effect. Gray: cis eQTLs without any significant *trans* effects.

**Figure S2-2.**
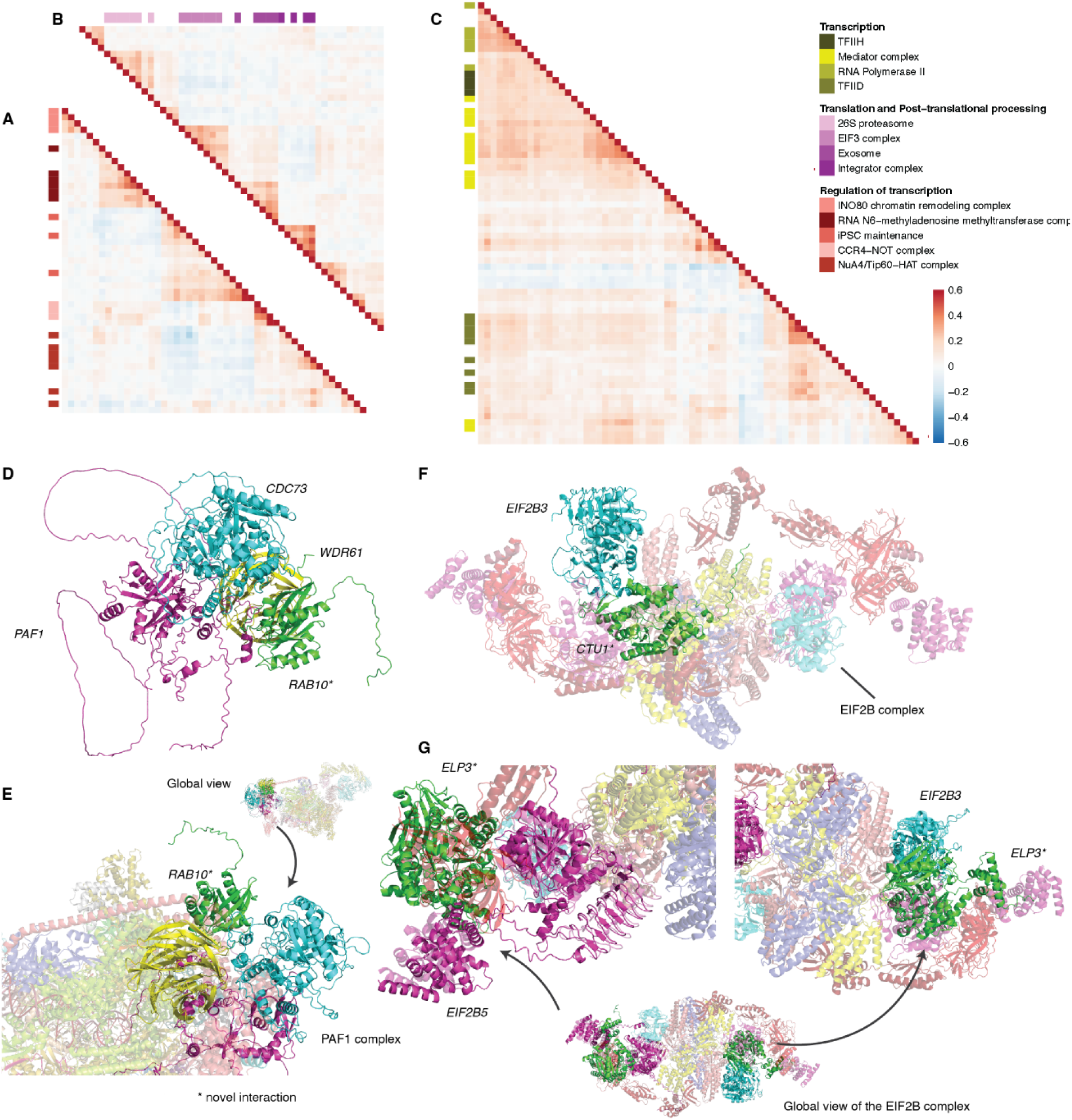
| Plausible protein interactions predicted from similar trans effects of knockdowns. Similarity of trans effects between target genes (x- and y-axis) involved in the A) regulation of transcription, B) translation and post-translational processing and C) transcription. Heatmap color: Pearson’s R. Annotation color: covariates of biological processes involved (see legend). D) A plausible quadramer formed between *RAB10*, *WDR61*, *CDC73* and *PAF1*. *RAB10* clashes with *CTR9* when we consider the larger structure of Paf^91^. E) Predicted binding structure between *RAB10* and the Paf complex. F) Predicted binding structure of *CTU1* with *EIF2B3* at a buried interface^92^ (6O81) of the EIF2B complex. G) Predicted binding structure of *ELP3* with *EIF2B3* and *EIF2B5* of the EIF2B complex.

**Figure S2-3.**
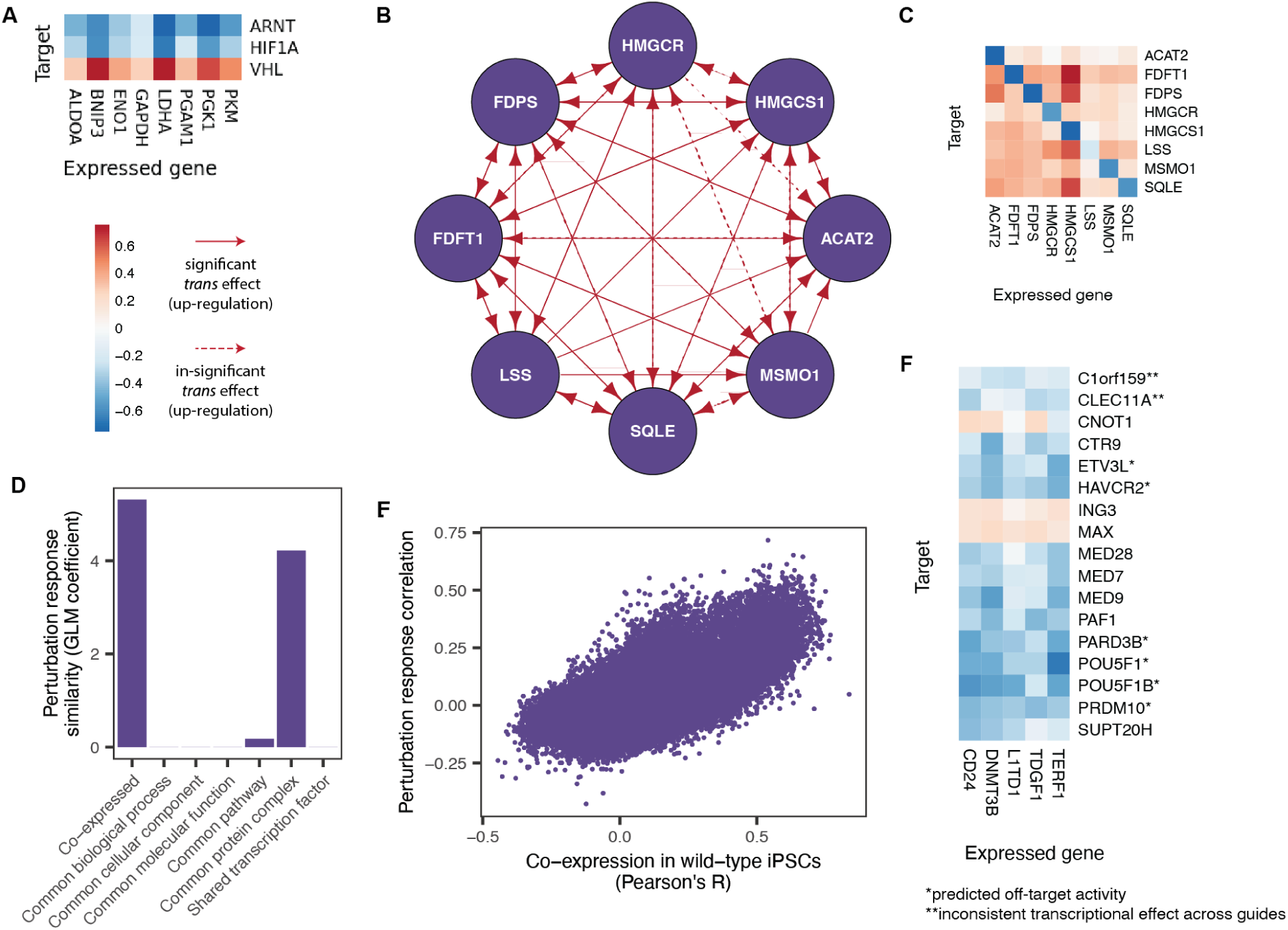
| Co-regulated modules identified by perturbation response similarity |. A) Heatmap of *trans* effects (log-fold change, color) of genes in the glycolysis pathway (x-axis) due to knockdown of hypoxia pathway regulators *ARNT*, *HIF1A* and *VHL* (y-axis). B) Joint up-regulation of cholesterol biosynthesis pathway members due to down-regulation of a pathway member. Purple nodes: Genes in the cholesterol biosynthesis pathway Red edges: up-regulation of arrow target upon knockdown of arrow source. C) As A), but change (color) of cholesterol biosynthesis gene expression (x-axis) upon knockdown of genes in cholesterol biosynthesis gene expression (y-axis). D) Predicting correlation between co-perturbation profiles of downstream effects. Coefficient (y-axis) for different covariates (x-axis) in a generalized linear model trained to predict correlation of downstream gene log-fold change vectors for pairs of targets. E) Correlation of gene expression values across single cells in wild-type iPSCs (*x*-axis) against correlation in response to perturbations of different targets in CRISPRi screening (*y*-axis). F) As A) and C) but of *trans* effects on iPSC marker genes *CD24*, *DNMT3B*, *L1TD1*, *TDGF1* and *TERF1* (x-axis) due to different target genes (y-axis). Single star: predicted off-target activity on *OCT4* (**Methods**). Double star: inconsistent transcriptional change between guides for the same gene (maximum correlation between guides < 0.1).

**Figure S3-1.**
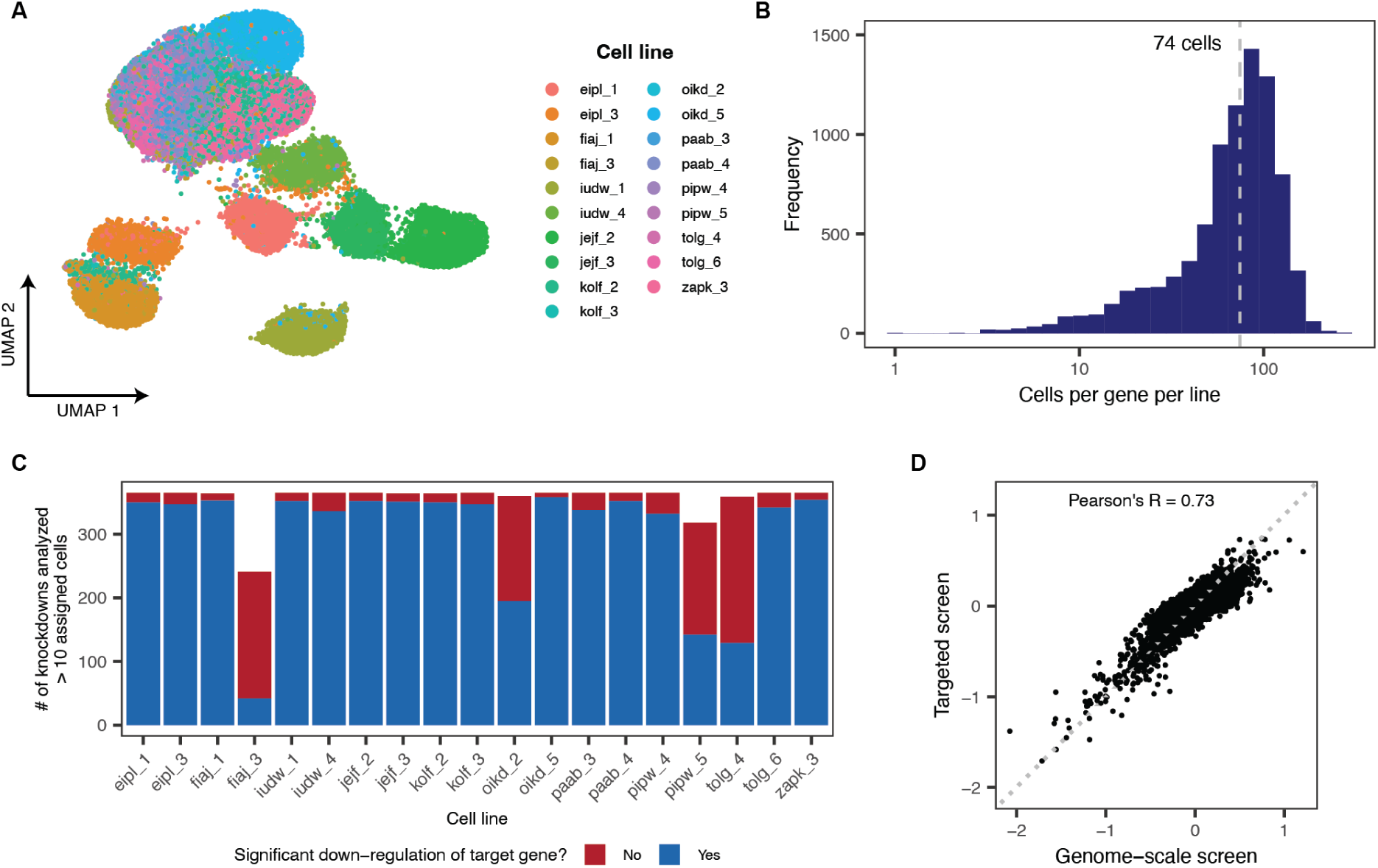
| Recovery of the targeted screen. A) UMAP representation (x- and y-axis) of technical covariate corrected expression of all assigned cells (markers) in the targeted screen, Colors: cell lines. B) Number of assigned cells (y-axis) per knockdown per line (x-axis). Dashed line: median number of cells per knockdown per line. C) Number of knockdowns with at least 10 assigned cells, plotted per line. Blue: number of knockdowns with significant (Benjamini-Hochberg adjusted p-value < 0.1, t-test) on-target down-regulation in a line. Red: additional number of knockdowns with insignificant on-target down-regulation. D) Concordance of all *trans* effects that were significant in either the genome-scale or target screen. Log-fold change across all cells in the targeted screen (y-axis) compared to genome-scale screen (x-axis) for 288,089 (target, downstream gene) pairs. A point represents a target-expressed gene pair.

**Figure S3-2.**
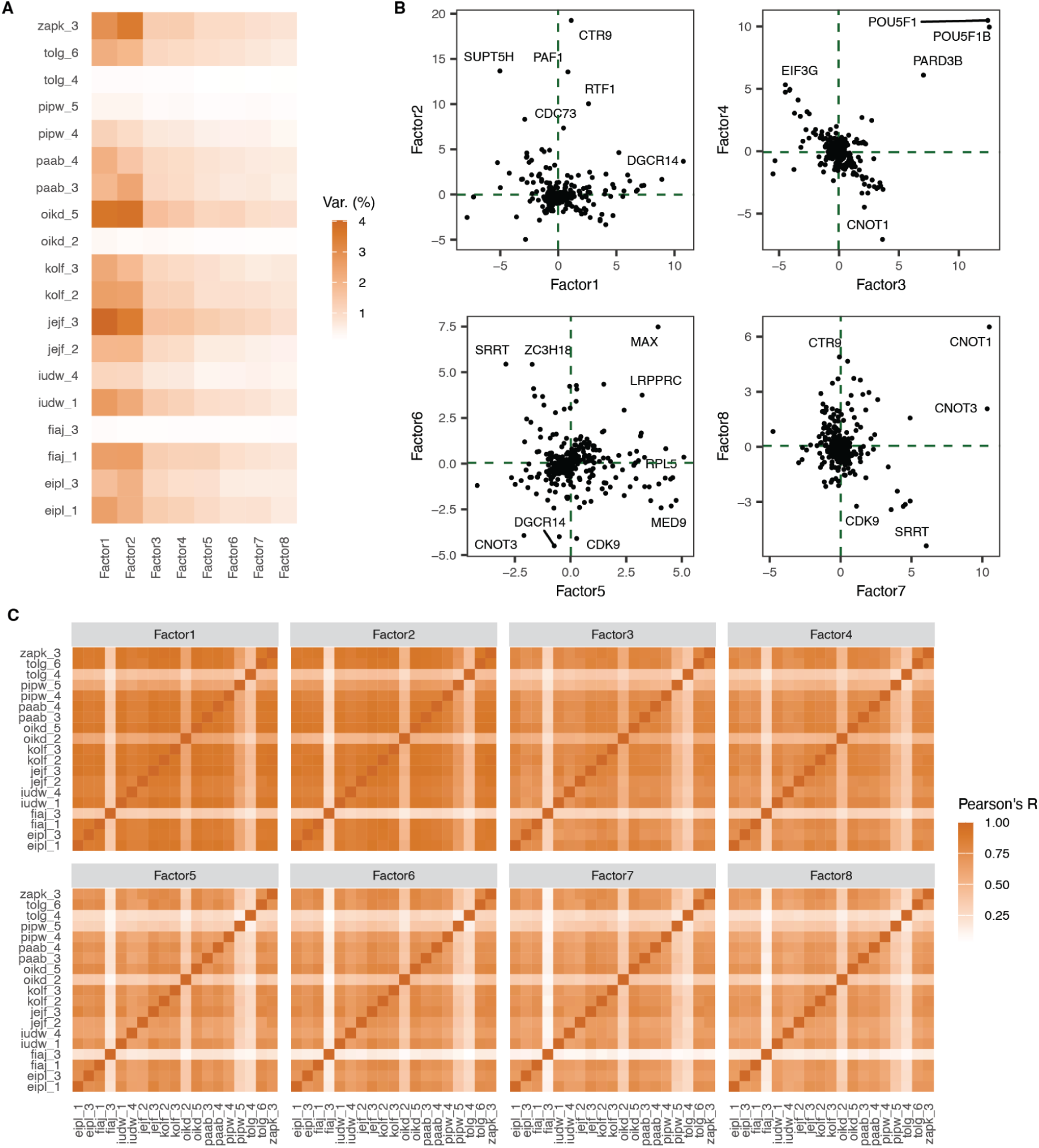
| Global effect of transcriptional change due CRISPRi perturbation across cell lines. A) Percentage of log-fold change variance explained (color) by different MOFA factors (x-axis) in different cell lines (y-axis). B) MOFA weights of a knockdown (markers) for different factors (x- and y- axis), for different factor combinations (panels). C) Correlation (color) of MOFA factors (panels), between cell lines (x- and y-axis).

**Figure S3-3.**
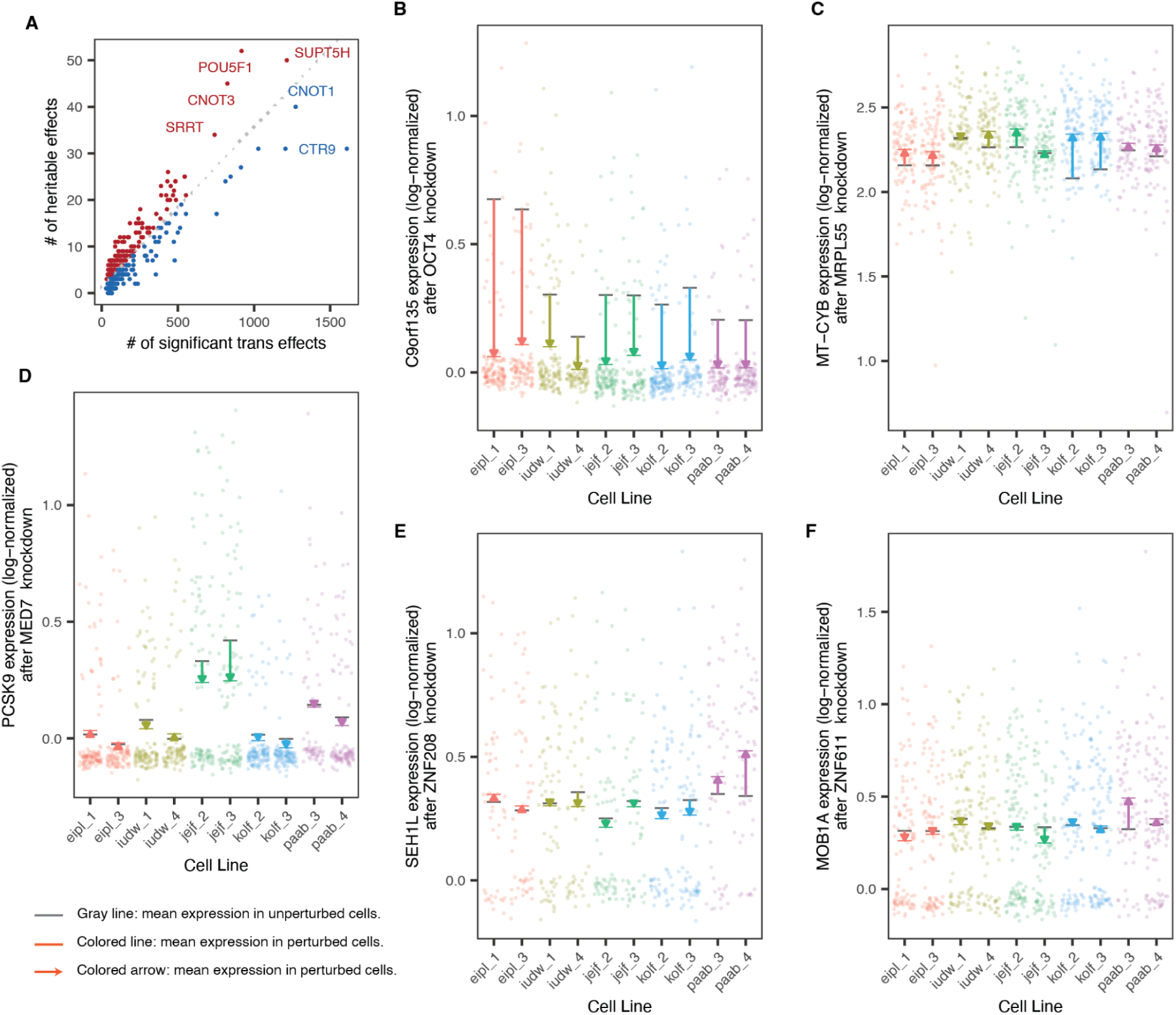
| Genetic background influences transcriptional response due to knockdown. A) Number of heritable *trans* effects vs. # of trans effects tested per gene. Genes in red indicate knockdowns with more heritable *trans* effects than expected, blue genes indicate knockdowns with fewer heritable *trans* effects than expected given the number of significant *trans* effects across lines. B) An example of loss of heritability. *C9orf135* expression change due to knockdown of *OCT4*. C9orf135 expression (y-axis; log-normalized) in individual cells (markers) from different cell lines (x-axis, colors) with *OCT4* knockdown. Colored dash: mean expression in knockdown in cell line. Grey dash: mean expression in control cells in cell line. Colored arrow: median expression change in line in response to knockdown. C) As B), but expression change of *MRPL55* due to knockdown of *MRPL55*. D) Expression change of *PCSK9* due to knockdown of *MED7*. E) Expression change of *SEH1L* due to knockdown of the *trans* eQTL hotspot *ZNF208*. F) Expression change of *MOB1A* due to knockdown of the *trans* eQTL hotspot *ZNF611*.

**Figure S4.**
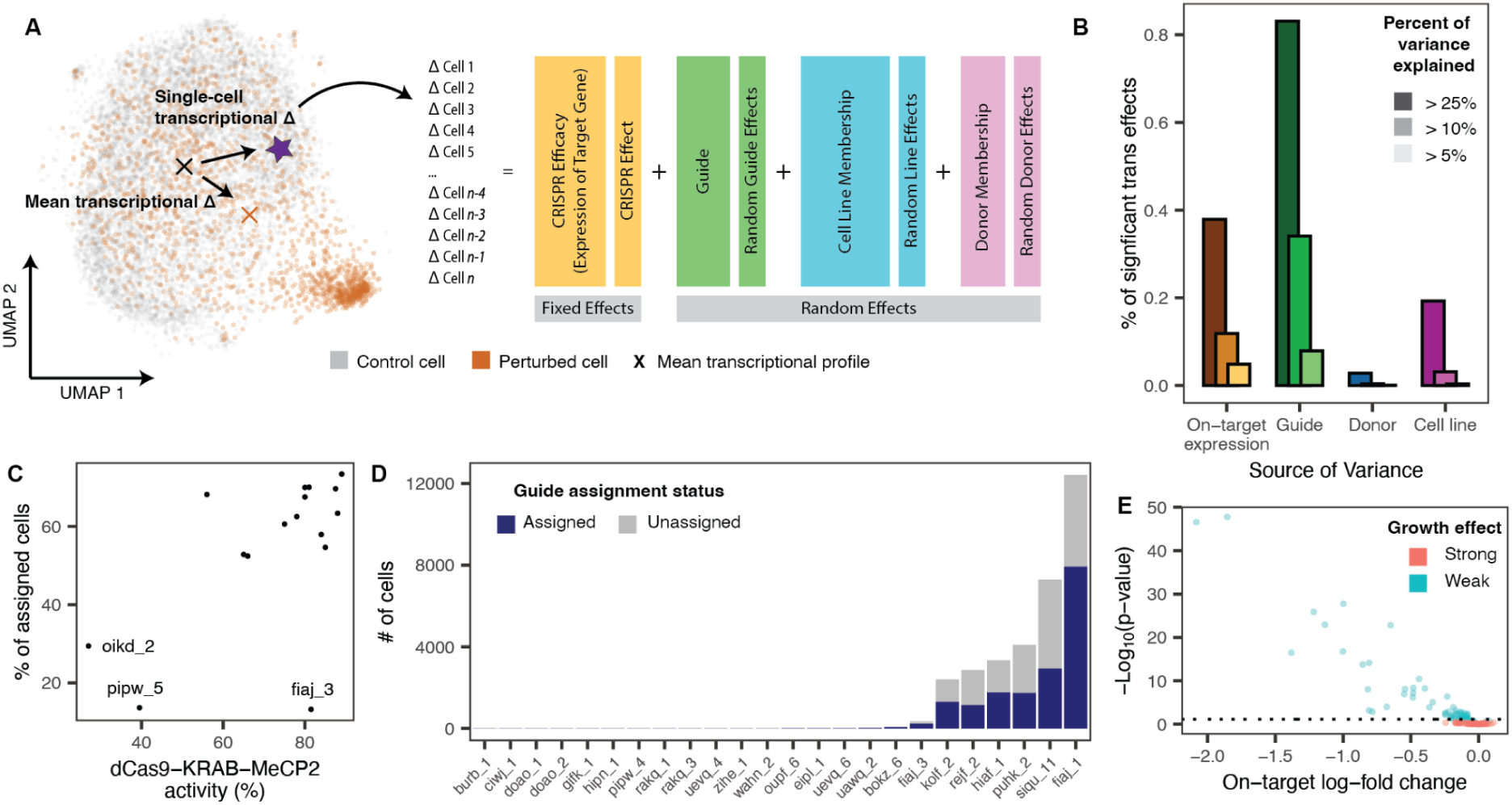
| Variation of transcriptional change due CRISPRi perturbation. A) Strategy for quantifying sources of variation in transcriptional response due to knockdown. B) Percentage of variance explained in transcriptional response due to knockdown due to CRISPRi efficacy (on-target expression), guide, donor and cell line. C) dCas9-KRAB-MeCP2 activity vs. fraction of assigned cells. D) Number of cells recovered per cell line after 14 days of selection (y-axis) for different cell lines (x-axis) in a pilot experiment with early pooling of lines. E) Repression log-fold change (x-axis) and log-scale p-value (y-axis) of target gene (markers) in a pilot experiment with early pooling of lines and late sequencing time point (14 days post-infection).

**Figure SM.**
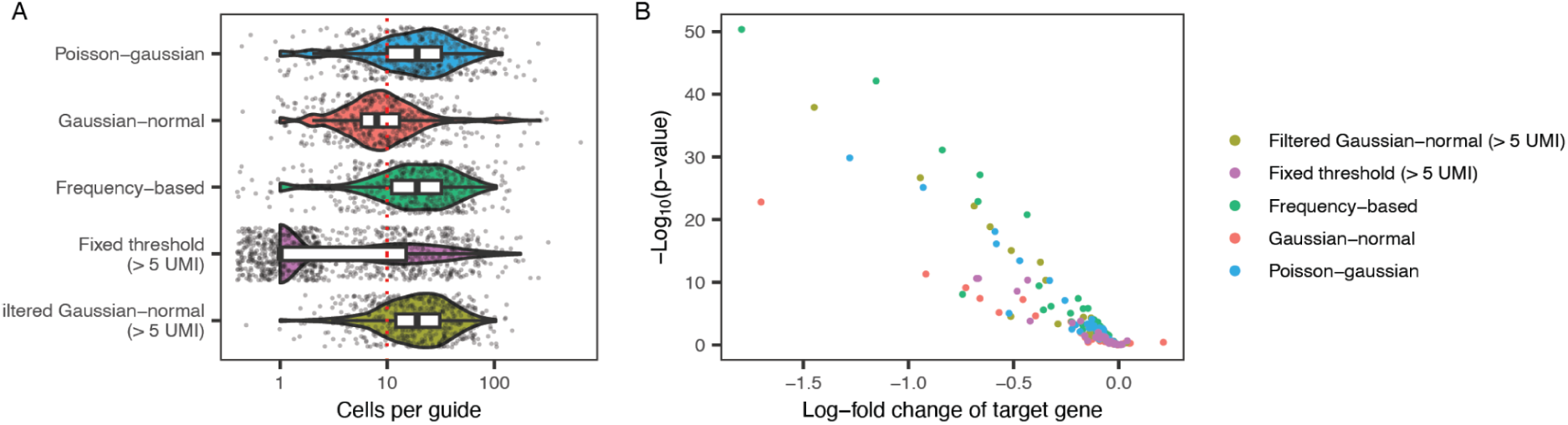
| Comparison of guide assignment strategies. We utilized a Poisson-Gaussian model^21^, a Gaussian-Gaussian model^69^ adopted from previous work, as well as assigning cells to a guide if and only if more than a fixed threshold (> 5) of guide UMIs were detected (fixed threshold), if and only if the fraction of a guide compared to all UMIs in a given cell was greater than a given threshold (ratio) and a modified version of the Gaussian-Gaussian model where assignments were further filtered so that assignments based on fewer than 5 guide UMIs were disregarded (filtered Gaussian-Gaussian). To evaluate the quality of each guide assignment method, we considered A) the number of assigned cells per guide and B) on-target repression.

