## Supplemental Table M-2 for "A genome-scale single cell CRISPRi map of *trans* gene regulation across human pluripotent stem cell lines"

| **Primer ID** | **Sequence** | **Subpool** |
| --- | --- | --- |
| 512 | GAGCACAGATGGCACAAATG | GenomeWideScreen_Fitness_1;  TargetedScreen |
| 513 | GAGCACAGATGGCAAACAGG | GenomeWideScreen_Fitness_2 |
| 514 | GAGCACAGATGGCACTTTTG | GenomeWideScreen_Fitness_3 |
| 515 | GAGCACAGATGGCTCTACGC | GenomeWideScreen_Fitness_4 |
| 516 | GAGCACAGATGGAGGACAGC | GenomeWideScreen_Fitness_5 |
| 517 | GAGCACAGATGGGTTGCTTG | GenomeWideScreen_Fitness_6 |
| 518 | GAGCACAGATGGACCTCGAG | GenomeWideScreen_Fitness_7 |
| 519 | GAGCACAGATGGCCTACGAG | GenomeWideScreen_Fitness_8 |
| 520 | GAGCACAGATGGTCATGTCG | GenomeWideScreen_Fitness_9 |
| 521 | GAGCACAGATGGCCTGGTAC | GenomeWideScreen_Fitness_10 |
| 522 | GAGCACAGATGGACTGGTTG | GenomeWideScreen_Fitness_11 |
| 523 | GAGCACAGATGGCGGTAAAG | GenomeWideScreen_Fitness_12 |
| 889 | ACAGCTCAGACGtcctgaga | GenomeWideScreen_Fitness_1  TargetedScreen |
| 890 | ACAGCTCAGACGggcttacc | GenomeWideScreen_Fitness_2 |
| 891 | ACAGCTCAGACGcaacggct | GenomeWideScreen_Fitness_3 |
| 892 | ACAGCTCAGACGgacggctc | GenomeWideScreen_Fitness_4 |
| 893 | ACAGCTCAGACGagccgtcg | GenomeWideScreen_Fitness_5 |
| 894 | ACAGCTCAGACGcatagcgc | GenomeWideScreen_Fitness_6 |
| 895 | ACAGCTCAGACGgaacatgc | GenomeWideScreen_Fitness_7 |
| 896 | ACAGCTCAGACGccgtacgt | GenomeWideScreen_Fitness_8 |
| 897 | ACAGCTCAGACGaacgcgtc | GenomeWideScreen_Fitness_9 |
| 898 | ACAGCTCAGACGacgttcag | GenomeWideScreen_Fitness_10 |
| 899 | ACAGCTCAGACGcctacgga | GenomeWideScreen_Fitness_11 |
| 900 | ACAGCTCAGACGtgactgcc | GenomeWideScreen_Fitness_12 |
| 1009 | agcagatctgcgaaatcggatccgc | NA |
| 1010 | TTCCTCTGCCCTCgccctcccacacataaccag | NA |
| 1012 | acaagttaacttatctagatccggtggatcccg | NA |
| 1013 | caccggatctagataagttaacttgtttattgcagctt | NA |
| 1014 | cgtgataacttattatatatatattttcttgttatagatatc | NA |
| 1016 | CATGCGGTGACGTGGAGGAGAATCCTGGCCCAATGGTGAGCAAGGGCGAGGC | NA |
