## Supplemental Table M-2 for "A genome-scale single cell CRISPRi map of *trans* gene regulation across human pluripotent stem cell lines"

**BFP**

ATGAGCGAGCTGATTAAGGAGAACATGCACATGAAGCTGTACATGGAGGGCACCGTGGAC

AACCATCACTTCAAGTGCACATCCGAGGGCGAAGGCAAGCCCTACGAGGGCACCCAGACC

ATGAGAATCAAGGTGGTCGAGGGCGGCCCTCTCCCCTTCGCCTTCGACATCCTGGCTACT

AGCTTCCTCTACGGCAGCAAGACCTTCATCAACCACACCCAGGGCATCCCCGACTTCTTC

AAGCAGTCCTTCCCTGAGGGCTTCACATGGGAGAGAGTCACCACATACGAGGACGGGGGC

GTGCTGACCGCTACCCAGGACACCAGCCTCCAGGACGGCTGCCTCATCTACAACGTCAAG

ATCAGAGGGGTGAACTTCACATCCAACGGCCCTGTGATGCAGAAGAAAACACTCGGCTGG

GAGGCCTTCACCGAGACGCTGTACCCCGCTGACGGCGGCCTGGAAGGCAGAAACGACATG

GCCCTGAAGCTCGTGGGCGGGAGCCATCTGATCGCAAACATCAAGACCACATATAGATCC

AAGAAACCCGCTAAGAACCTCAAGATGCCTGGCGTCTACTATGTGGACTACAGACTGGAA

AGAATCAAGGAGGCCAACAACGAGACCTACGTCGAGCAGCACGAGGTGGCAGTGGCCAGA

TACTGCGACCTCCCTAGCAAACTGGGGCACAAGCTTAAT

**BSD**

ATGGCCAAGCCTTTGTCTCAAGAAGAATCCACCCTCATTGAAAGAGCAACGGCTACAATC

AACAGCATCCCCATCTCTGAAGACTACAGCGTCGCCAGCGCAGCTCTCTCTAGCGACGGC

CGCATCTTCACTGGTGTCAATGTATATCATTTTACTGGGGGACCTTGTGCAGAACTCGTG

GTGCTGGGCACTGCTGCTGCTGCGGCAGCTGGCAACCTGACTTGTATCGTCGCGATCGGA

AATGAGAACAGGGGCATCTTGAGCCCCTGCGGACGGTGCCGACAGGTGCTTCTCGATCTG

CATCCTGGGATCAAAGCCATAGTGAAGGACAGTGATGGACAGCCGACGGCAGTTGGGATT

CGTGAATTGCTGCCCTCTGGTTATGTGTGGGAGGGC

**dCas9-KRAB-MeCP2**

ATGGACAAGAAGTACTCCATTGGGCTCGCTATCGGCACAAACAGCGTCGGCTGGGCCGTC

ATTACGGACGAGTACAAGGTGCCGAGCAAAAAATTCAAAGTTCTGGGCAATACCGATCGC

CACAGCATAAAGAAGAACCTCATTGGCGCCCTCCTGTTCGACTCCGGGGAGACGGCCGAA

GCCACGCGGCTCAAAAGAACAGCACGGCGCAGATATACCCGCAGAAAGAATCGGATCTGC

TACCTGCAGGAGATCTTTAGTAATGAGATGGCTAAGGTGGATGACTCTTTCTTCCATAGG

CTGGAGGAGTCCTTTTTGGTGGAGGAGGATAAAAAGCACGAGCGCCACCCAATCTTTGGC

AATATCGTGGACGAGGTGGCGTACCATGAAAAGTACCCAACCATATATCATCTGAGGAAG

AAGCTTGTAGACAGTACTGATAAGGCTGACTTGCGGTTGATCTATCTCGCGCTGGCGCAT

ATGATCAAATTTCGGGGACACTTCCTCATCGAGGGGGACCTGAACCCAGACAACAGCGAT

GTCGACAAACTCTTTATCCAACTGGTTCAGACTTACAATCAGCTTTTCGAAGAGAACCCG

ATCAACGCATCCGGAGTTGACGCCAAAGCAATCCTGAGCGCTAGGCTGTCCAAATCCCGG

CGGCTCGAAAACCTCATCGCACAGCTCCCTGGGGAGAAGAAGAACGGCCTGTTTGGTAAT

CTTATCGCCCTGTCACTCGGGCTGACCCCCAACTTTAAATCTAACTTCGACCTGGCCGAA

GATGCCAAGCTTCAACTGAGCAAAGACACCTACGATGATGATCTCGACAATCTGCTGGCC

CAGATCGGCGACCAGTACGCAGACCTTTTTTTGGCGGCAAAGAACCTGTCAGACGCCATT

CTGCTGAGTGATATTCTGCGAGTGAACACGGAGATCACCAAAGCTCCGCTGAGCGCTAGT

ATGATCAAGCGCTATGATGAGCACCACCAAGACTTGACTTTGCTGAAGGCCCTTGTCAGA

CAGCAACTGCCTGAGAAGTACAAGGAAATTTTCTTCGATCAGTCTAAAAATGGCTACGCC

GGATACATTGACGGCGGAGCAAGCCAGGAGGAATTTTACAAATTTATTAAGCCCATCTTG

GAAAAAATGGACGGCACCGAGGAGCTGCTGGTAAAGCTTAACAGAGAAGATCTGTTGCGC

AAACAGCGCACTTTCGACAATGGAAGCATCCCCCACCAGATTCACCTGGGCGAACTGCAC

GCTATCCTCAGGCGGCAAGAGGATTTCTACCCCTTTTTGAAAGATAACAGGGAAAAGATT

GAGAAAATCCTCACATTTCGGATACCCTACTATGTAGGCCCCCTCGCCCGGGGAAATTCC

AGATTCGCGTGGATGACTCGCAAATCAGAAGAGACCATCACTCCCTGGAACTTCGAGGAA

GTCGTGGATAAGGGGGCCTCTGCCCAGTCCTTCATCGAAAGGATGACTAACTTTGATAAA

AATCTGCCTAACGAAAAGGTGCTTCCTAAACACTCTCTGCTGTACGAGTACTTCACAGTT

TATAACGAGCTCACCAAGGTCAAATACGTCACAGAAGGGATGAGAAAGCCAGCATTCCTG

TCTGGAGAGCAGAAGAAAGCTATCGTGGACCTCCTCTTCAAGACGAACCGGAAAGTTACC

GTGAAACAGCTCAAAGAAGACTATTTCAAAAAGATTGAATGTTTCGACTCTGTTGAAATC

AGCGGAGTGGAGGATCGCTTCAACGCATCCCTGGGAACGTATCACGATCTCCTGAAAATC

ATTAAAGACAAGGACTTCCTGGACAATGAGGAGAACGAGGACATTCTTGAGGACATTGTC

CTCACCCTTACGTTGTTTGAAGATAGGGAGATGATTGAAGAACGCTTGAAAACTTACGCT

CATCTCTTCGACGACAAAGTCATGAAACAGCTCAAGAGGCGCCGATATACAGGATGGGGG

CGGCTGTCAAGAAAACTGATCAATGGGATCCGAGACAAGCAGAGTGGAAAGACAATCCTG

GATTTTCTTAAGTCCGATGGATTTGCCAACCGGAACTTCATGCAGTTGATCCATGATGAC

TCTCTCACCTTTAAGGAGGACATCCAGAAAGCACAAGTTTCTGGCCAGGGGGACAGTCTT

CACGAGCACATCGCTAATCTTGCAGGTAGCCCAGCTATCAAAAAGGGAATACTGCAGACC

GTTAAGGTCGTGGATGAACTCGTCAAAGTAATGGGAAGGCATAAGCCCGAGAATATCGTT

ATCGAGATGGCCCGAGAGAACCAAACTACCCAGAAGGGACAGAAGAACAGTAGGGAAAGG

ATGAAGAGGATTGAAGAGGGTATAAAAGAACTGGGGTCCCAAATCCTTAAGGAACACCCA

GTTGAAAACACCCAGCTTCAGAATGAGAAGCTCTACCTGTACTACCTGCAGAACGGCAGG

GACATGTACGTGGATCAGGAACTGGACATCAATCGGCTCTCCGACTACGACGTGGCTGCT

ATCGTGCCCCAGTCTTTTCTCAAAGATGATTCTATTGATAATAAAGTGTTGACAAGATCC

GATAAAGCTAGAGGGAAGAGTGATAACGTCCCCTCAGAAGAAGTTGTCAAGAAAATGAAA

AATTATTGGCGGCAGCTGCTGAACGCCAAACTGATCACACAACGGAAGTTCGATAATCTG

ACTAAGGCTGAACGAGGTGGCCTGTCTGAGTTGGATAAAGCCGGCTTCATCAAAAGGCAG

CTTGTTGAGACACGCCAGATCACCAAGCACGTGGCCCAAATTCTCGATTCACGCATGAAC

ACCAAGTACGATGAAAATGACAAACTGATTCGAGAGGTGAAAGTTATTACTCTGAAGTCT

AAGCTGGTCTCAGATTTCAGAAAGGACTTTCAGTTTTATAAGGTGAGAGAGATCAACAAT

TACCACCATGCGCATGATGCCTACCTGAATGCAGTGGTAGGCACTGCACTTATCAAAAAA

TATCCCAAGCTTGAATCTGAATTTGTTTACGGAGACTATAAAGTGTACGATGTTAGGAAA

ATGATCGCAAAGTCTGAGCAGGAAATAGGCAAGGCCACCGCTAAGTACTTCTTTTACAGC

AATATTATGAATTTTTTCAAGACCGAGATTACACTGGCCAATGGAGAGATTCGGAAGCGA

CCACTTATCGAAACAAACGGAGAAACAGGAGAAATCGTGTGGGACAAGGGTAGGGATTTC

GCGACAGTCCGGAAGGTCCTGTCCATGCCGCAGGTGAACATCGTTAAAAAGACCGAAGTA

CAGACCGGAGGCTTCTCCAAGGAAAGTATCCTCCCGAAAAGGAACAGCGACAAGCTGATC

GCACGCAAAAAAGATTGGGACCCCAAGAAATACGGCGGATTCGATTCTCCTACAGTCGCT

TACAGTGTACTGGTTGTGGCCAAAGTGGAGAAAGGGAAGTCTAAAAAACTCAAAAGCGTC

AAGGAACTGCTGGGCATCACAATCATGGAGCGATCAAGCTTCGAAAAAAACCCCATCGAC

TTTCTCGAGGCGAAAGGATATAAAGAGGTCAAAAAAGACCTCATCATTAAGCTTCCCAAG

TACTCTCTCTTTGAGCTTGAAAACGGCCGGAAACGAATGCTCGCTAGTGCGGGCGAGCTG

CAGAAAGGTAACGAGCTGGCACTGCCCTCTAAATACGTTAATTTCTTGTATCTGGCCAGC

CACTATGAAAAGCTCAAAGGGTCTCCCGAAGATAATGAGCAGAAGCAGCTGTTCGTGGAA

CAACACAAACACTACCTTGATGAGATCATCGAGCAAATAAGCGAATTCTCCAAAAGAGTG

ATCCTCGCCGACGCTAACCTCGATAAGGTGCTTTCTGCTTACAATAAGCACAGGGATAAG

CCCATCAGGGAGCAGGCAGAAAACATTATCCACTTGTTTACTCTGACCAACTTGGGCGCG

CCTGCAGCCTTCAAGTACTTCGACACCACCATAGACAGAAAGCGGTACACCTCTACAAAG

GAGGTCCTGGACGCCACACTGATTCATCAGTCAATTACGGGGCTCTATGAAACAAGAATC

GACCTCTCTCAGCTCGGTGGAGAC

**mScarlet**

ATGCGGTTCAAGGTGCACATGGAGGGCTCCATGAACGGCCACGAGTTCGAGATCGAGGGC

GAGGGCGAGGGCCGCCCCTACGAGGGCACCCAGACCGCCAAGCTGAAGGTGACCAAGGGT

GGCCCCCTGCCCTTCTCCTGGGACATCCTGTCCCCTCAGTTCATGTACGGCTCCAGGGCC

TTCACCAAGCACCCCGCCGACATCCCCGACTACTATAAGCAGTCCTTCCCCGAGGGCTTC

AAGTGGGAGCGCGTGATGAACTTCGAGGACGGCGGCGCCGTGACCGTGACCCAGGACACC

TCCCTGGAGGACGGCACCCTGATCTACAAGGTGAAGCTCCGCGGCACCAACTTCCCTCCT

GACGGCCCCGTAATGCAGAAGAAGACAATGGGCTGGGAAGCGTCCACCGAGCGGTTGTAC

CCCGAGGACGGCGTGCTGAAGGGCGACATTAAGATGGCCCTGCGCCTGAAGGACGGCGGC

CGCTACCTGGCGGACTTCAAGACCACCTACAAGGCCAAGAAGCCCGTGCAGATGCCCGGC

GCCTACAACGTCGACCGCAAGTTGGACATCACCTCCCACAACGAGGACTACACCGTGGTG

GAACAGTACGAACGCTCCGAGGGCCGCCACTCCACCGGCGGCATGGACGAGCTGTACAAG

TCCGGACTCAGATCTCGAGCTCAAGCTTCGAATTCTGCAGTCGACGGTACCGCGGGCCCG

GGATCCACCGGATCTAGA

**PURO**

ATGACCGAGTACAAGCCCACGGTGCGCCTCGCCACCCGCGACGACGTCCCCAGGGCCGTA

CGCACCCTCGCCGCCGCGTTCGCCGACTACCCCGCCACGCGCCACACCGTCGATCCGGAC

CGCCACATCGAGCGGGTCACCGAGCTGCAAGAACTCTTCCTCACGCGCGTCGGGCTCGAC

ATCGGCAAGGTGTGGGTCGCGGACGACGGCGCCGCGGTGGCGGTCTGGACCACGCCGGAG

AGCGTCGAAGCGGGGGCGGTGTTCGCCGAGATCGGCCCGCGCATGGCCGAGTTGAGCGGT

TCCCGGCTGGCCGCGCAGCAACAGATGGAAGGCCTCCTGGCGCCGCACCGGCCCAAGGAG

CCCGCGTGGTTCCTGGCCACCGTCGGCGTCTCGCCCGACCACCAGGGCAAGGGTCTGGGC

AGCGCCGTCGTGCTCCCCGGAGTGGAGGCGGCCGAGCGCGCCGGGGTGCCCGCCTTCCTG

GAGACCTCCGCGCCCCGCAACCTCCCCTTCTACGAGCGGCTCGGCTTCACCGTCACCGCC

GACGTCGAGGTGCCCGAAGGACCGCGCACCTGGTGCATGACCCGCAAGCCCGGTGCC
